# Impairment in O-acetylserine-(thiol) lyase A and B, but not C, confers higher selenate sensitivity and uncovers role for A, B and C as L-Cys and L-SeCys desulfhydrases in *Arabidopsis*

**DOI:** 10.1101/2020.09.16.300020

**Authors:** Assylay Kurmanbayeva, Aizat Bekturova, Aigerim Soltabayeva, Sudhakar Srivastava, Dinara Oshanova, Zhadyrassyn Nurbekova, Moshe Sagi

## Abstract

The role of the cytosolic O-acetylserine-(thiol) lyase A (OASTLA), chloroplastic OASTLB and mitochondrion OASTLC in plant resistance/sensitivity to selenate was studied in *Arabidopsis* plants. Impairment in OASTLA and B resulted in reduced biomass, chlorophyll and soluble protein levels compared with impaired OASTL C and Wild-Type treated with selenate. The lower organic-Se and protein-Se levels followed by decreased organic-S, S in proteins and total glutathione in *oastlA* and *oastlB* compared to Wild-Type and *oastlC* are indicative that Se accumulation is not the main cause for the stress symptoms, but rather the interference of Se with the S-reduction pathway. The increase in sulfite oxidase, adenosine 5′-phosphosulfate reductase, sulfite reductase and OASTL activity levels, followed by enhanced sulfite and sulfide, indicate a futile anabolic S-starvation response to selenate-induced organic-S catabolism in *oastlA* and *oastlB* compared to Wild-Type and *oastlC*.

Additionally, the catabolic pathway of L-cysteine degradation was enhanced by selenate, and similar to L-cysteine producing activity, *oastlA* and *B* exhibited a significant decrease in L-cysteine desulfhydrase (DES) activity, compared with WT, indicating a major role of OASTLs in L-cysteine degradation. This notion was further evidenced by sulfide dependent DES in-gel activity, immunoblotting, immunoprecipitation with specific antibodies and identification of unique peptides in activity bands generated by OASTLA, B and C. Similar responses of the OASTLs in Seleno-Cysteine degradation was demonstrated in selenate stressed plants. Notably, no L-cysteine and L-Seleno-Cysteine DES activity bands but those related to OASTLs were evident. These results indicate the significance of OASTLs in degrading L-cysteine and L-SelenoCysteine in *Arabidopsis*.

**Summary:** The cytosolic OASTLA and chloroplastic OASTLB have significantly higher desulfhydrase activity rates than the cytosolic DES1 and are able to degrade L-Cys and L-SeCys to sulfide and selenide, respectively in *Arabidopsis*.

## INTRODUCTION

The amino acid selenocysteine, a component of selenoproteins in humans, contains Se (Papp et al., 2007). Plants serve as a source of Se for mammalians; yet Se is not essential for plants. The chemical similarity of Se to S, enables plants to readily uptake Se via root sulfate transporters (Pilon-Smits and Quinn, 2010; Sors et al., 2005) and incorporate it into organic Se compounds via the S assimilation pathway components (Grant et al., 2011). The pathway (see Supplemental Fig. S1) is initiated with the adenylation of sulfate/selenate by ATP sulfurylase (ATPS, EC 2.7.7.4) to generate adenosine 5′-phosphosulfate (APS)/adenosine 5′-phosphoselenate (APSe). APS/APSe is then reduced to sulfite/selenite by the plastidic APS reductase (APR, EC 1.8.99.2) (Mroczek-Zdyrska and Wójcik, 2012; Schiavon et al., 2015; White, 2015). The toxic sulfite/selenite is further reduced to sulfide by the chloroplast-localized sulfite reductase (SiR, EC 1.8.7.1) (Khan et al., 2010; Yarmolinsky et al., 2013), whereas selenite can be reduced enzymatically to selenide by SiR and non-enzymatically by reduced glutathione [GSH (Fisher et al., 2016; Seko et al., 1989; White, 2015)]. Further, the generated sulfide/selenide together with O-acetylserine (OAS), the latter catalyzed by serine acetyltransferase (SAT, EC 2.3.1.30), are incorporated into cysteine (Cys) or selenocysteine (SeCys) in a reaction catalyzed by the O-acetylserine-(thiol) lyase (OASTL, EC 2.5.1.47) in cytosol (OASTLA), plastid (OASTLB), and mitochondria (OASTLC) [(Pilon-Smits and Womg, 2012; White, 2015; Wirtz and Hell, 2006) (see in Supplemental Fig. S1)]. The replacement of Cys by SeCys in proteins is thought to be a cause for selenate toxicity in *Arabidopsis* (Sabbagh and Van Hoewyk, 2012; Van Hoewyk, 2018).

Three major Se toxicity mechanisms were recently concluded in plants: competition between Se and S in primary and secondary metabolism, incorporation of SeCys and SeMet into proteins, and disruption caused by oxidative stress of metabolism and cellular structures (Lima et al., 2018; White, 2018). Amongst others, the role of Cys catabolic enzymes in L-SeCys degradation was not shown. Moreover, while the cytosolic L-cysteine desulfhydrase1 (DES1, EC 4.4.1.1) was shown to catalyze the degradation of cysteine to sulfide, pyruvate and ammonia (Alvarez et al., 2012b; Gotor et al., 2013; Romero et al., 2013), DES1 or other L-Cys catabolic enzymes were not shown to increase L-Cys and/or L-SeCys degradation in response to selenate.

In the current study we show that *oastlA and oastlB Atabidopsis* mutants are more sensitive to the presence of selenate in the growth medium than *oastlC* and WT. The significantly lower selenium level in the protein fraction and biomass of *oastlA* and *oastlB* compared to WT and *oastlC* mutant led us to explore the interference of selenate in the sulfate reduction pathway as the cause for the lower growth rate in the two mutants. The results demonstrate that in the absence of active OASTLA and OASTLB, the selenate interference in the S assimilation pathway leads to a decrease in organic S, S in protein fraction and total glutathione and has a more negative effect on the resulting biomass accumulation than the expected positive effect of lower selenium content in the organic fraction. Increase in the activity of core enzymes of the sulfate reduction pathway, the APR and SiR, followed by enhanced sulfite and sulfide levels uncovers a futile anabolic S-starvation response to the selenate induced organic-S catabolism in *oastlA* and *oastlB* compared to WT and *oastlC*. A major role for the OASTL proteins in acting as L-Cys and L-SeCys desulfhydrases in control and Selenate stressed plants was demonstrated as well.

## RESULTS

### *oastlA* and *oastlB* mutants are more sensitive to selenate than *WT* and *oastlC*

Replacement of cysteine by selenocysteine in proteins is considered to be toxic to plants (Lima et al., 2018; White, 2018); whereas minimizing the incorporation of selenocysteine into proteins has been shown to be an effective strategy to increase Se tolerance in plants (Van Hoewyk et al., 2005). *Arabidopsis* mutants impaired in the OASTLs: the cytosolic (*oastlA*), plastidal (*oastlB*) and mitochondrial (*oastlC*), were employed to examine whether these impairments will reduce selenium in the protein fraction and thus enhance plant tolerance to selenium toxicity.

The mutation of the various OASTL mutant plants was verified by immunoblot analysis, using an OASTL C polyclonal antiserum which also cross-reacts with OASTL A and OASTL B (kindly provided by Prof. Dr. S. Kopriva, University of Cologne, Köln). The three bands detected in WT fractionated proteins corresponded to OASTL A, B, and C from bottom to top respectively, whereas the lower, middle or upper bands were missing in the fractionated proteins of *oastlA, oastlB* or *oastlC* mutants respectively (Supplemental Fig. S2A).

*oastlA, oastlB* or *oastlC* mutants grown with *WT* plants on 0.5 MS agar medium in the presence and absence (control) of 40 µM selenate for 2 weeks exhibited similar growth as *WT* plant in the absence of selenate and showed a significant reduction in growth rate and chlorophyll level in response to selenate. Yet, the reduction in biomass and chlorophyll content shown in *oastlA* and *oastlB* mutants was significantly higher than in *WT* and *oastlC* (Fig. 1A-D), in spite of the lower Se level in the protein fraction of the *oastlA* and *oastlB* (Fig. 1E).

**Figure 1.**
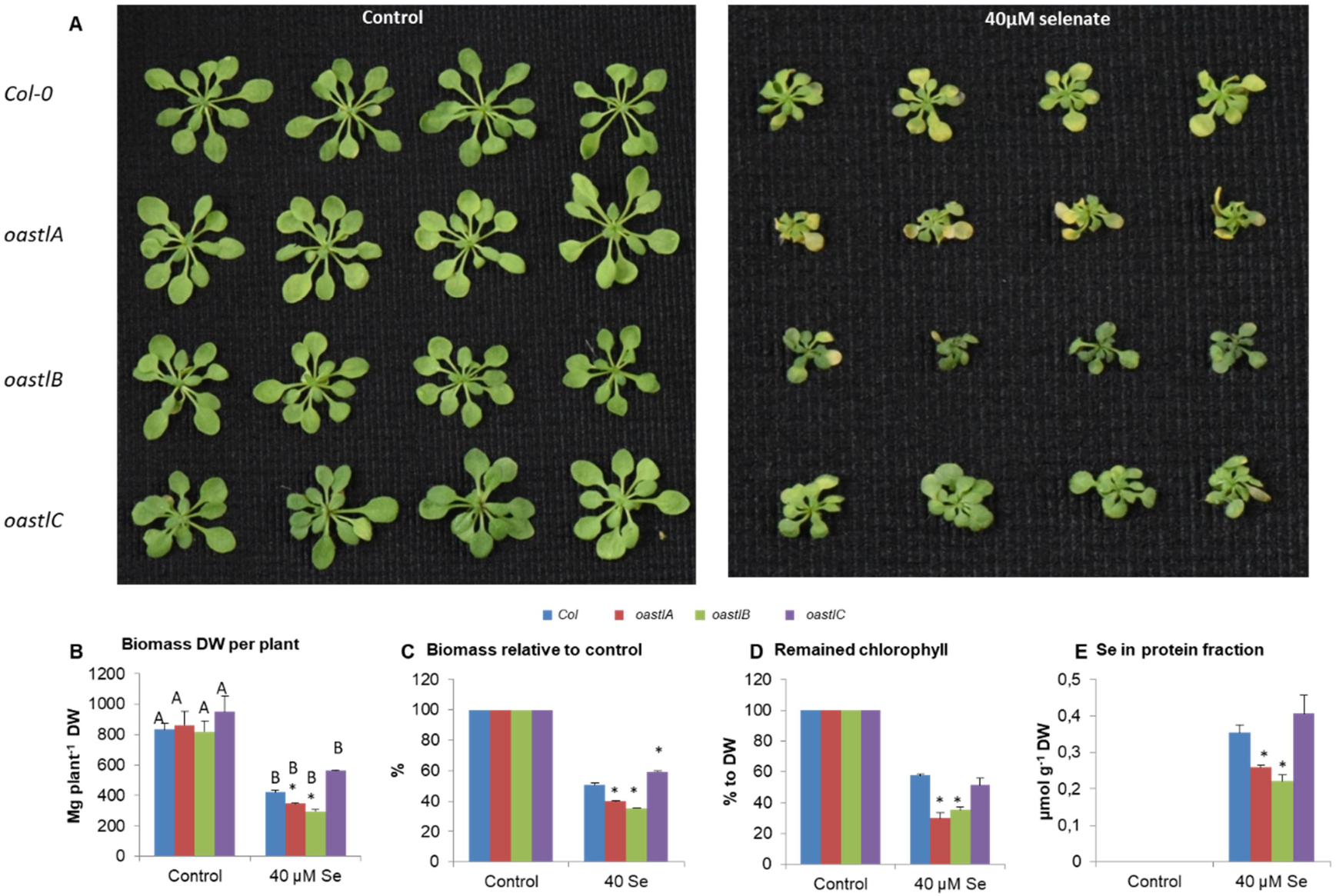
The effect of 40 µM selenate treatment on growth parameters and Se in protein fraction in WT, *oastlA, oastlB* and *oastlC KO* mutants of *Arabidopsis thaliana*. A. Phenotype of plants grown for 14 days under control (left panel) and 40µM selenate treated (right panel) conditions. The weight of plant biomass accumulation (B) and relative weight to their own controls (C) supplemented with or without selenate for 14 days. The values are means ± SE (n = 15). Remaining chlorophyll level (D) and Se in protein fraction (E). The values are means ± SE (n = 4). Values denoted with different letters are significantly different according to the T-test analyses, P < 0.05 (JMP 8.0 software, http://www.jmp.com/). Different uppercase letters indicate significant differences between control and Se treatment of each genotype. Asterisks indicate significant differences between *WT* and *oas-tlA, oas-tlB, oastlC* KO plants subjected to the same treatment.

Since at high levels selenate and Se can act as pro-oxidants and cause oxidative stress (Grant et al., 2011; Mroczek-Zdyrska and Wójcik, 2012), Se and selenate levels were determined in the leaves. Yet, no higher total Se and selenate were detected in *oastlA* and *oastlB* as compared to *WT* and *oastlC* (Supplemental Fig. S2B and C) which could be attributed to the decreased biomass accumulation and higher chlorophyll degradation rates in *oastlA* and *oastlB* mutants. These results indicate that the levels of Se in the protein fraction, total Se and selenate in leaf tissue are not the only cause for the higher sensitivity to Selenate noticed in *oastlA* and *oastlB* compared to WT and *oastlC* mutant.

### The interference in sulfate assimilation by selenate is stronger in *oastA* and *B* than in WT and *oastlC* mutant

Since higher sensitivity in the *oastlA* and *oastlB* mutants was not related to Se content in the leaves, the sulfate reduction pathway was further studied, estimating the level of selenate interference. Cysteine biosynthesis, the last step of the sulfate reduction pathway catalyzed by OASTLs was studied in the three mutants in comparison to WT. *OASTLA, OASTLB* or *OASTLC* transcripts were not expressed in *oastlA, oastlB* or *oastlC* mutants respectively, indicating the complete knockout of these genes (Supplemental Fig. S2D-F). Importantly, under control conditions *OASTLB* transcript was enhanced in *AtoastA* and *C* mutants, *OASTLA* in *AtoastB*, whereas *OASTC* was enhanced in *AtoastB*, indicating a level of complementation to compensate for the mutation among the various transcripts. The presence of selenate in the growth medium resulted in a significant enhancement of the transcripts, exhibiting the increase of the three transcripts in *WT*, as well as *OASTLA* and *B* transcripts in *oastlC*, whereas in *oastlA* the transcript of *OASTLB* and in *oastlB OASTLA*, both were significantly increased (Supplemental Fig. S2D-F). Similarly, the absence of either OASTLA or C in *oastlA* or *oastlC* mutants respectively, resulted in the enhancement of the two other functioning OASTLs in each of the mutants as compared to the proteins expression in WT leaves grown under unstressed conditions, whereas in *oastlB* only OASTLA exhibited such response (Supplemental Fig. S2A). Supplementation of selenate to the growth medium resulted in increased expression of OASTLB and C proteins in WT leaves, and the enhancement of OASTLB in *oastlA*, OASTLA in *oastlB* and OASTLB in *AtoastlC* as compared to their control unstressed plants (Supplemental Fig. S2A).

In-spite of the enhanced protein expression of OASTLB and C in the absence of selenate and of OASTLB in the presence of selenate, the impairment in OASTLA resulted in a strongly decreased cysteine production activity of OASTL in *AtoastlA* mutant, exhibiting a reduction in activity by 87% and 72% as compared to WT under control and selenate treated conditions, respectively. The *AtoastlB* mutant showed a significant decline of 28% in both conditions, while in *oastlC* OASTL activity was similar to the activity in WT plants. Interestingly, the activity in all the examined genotypes was significantly increased with selenate treatment, likely as the result of the enhanced expression of at least one of the active OASTL proteins (Fig. 2A). The comparable activity rates to WT found in *oastlC* mutant might be the result of the enhanced expression of the two major proteins, the OASTLA and B as compared to WT in control as well as under stressed conditions (Supplemental Fig. S2A). Taken together, the enhanced OASTL activity in leaves of selenate stressed WT and the OASTL mutants may indicate S-limitation type response shown before in *Arabidopsis* leaves in response to limited S supply (Barraso et al., 1995; Hesse et al., 1999).

**Figure 2.**
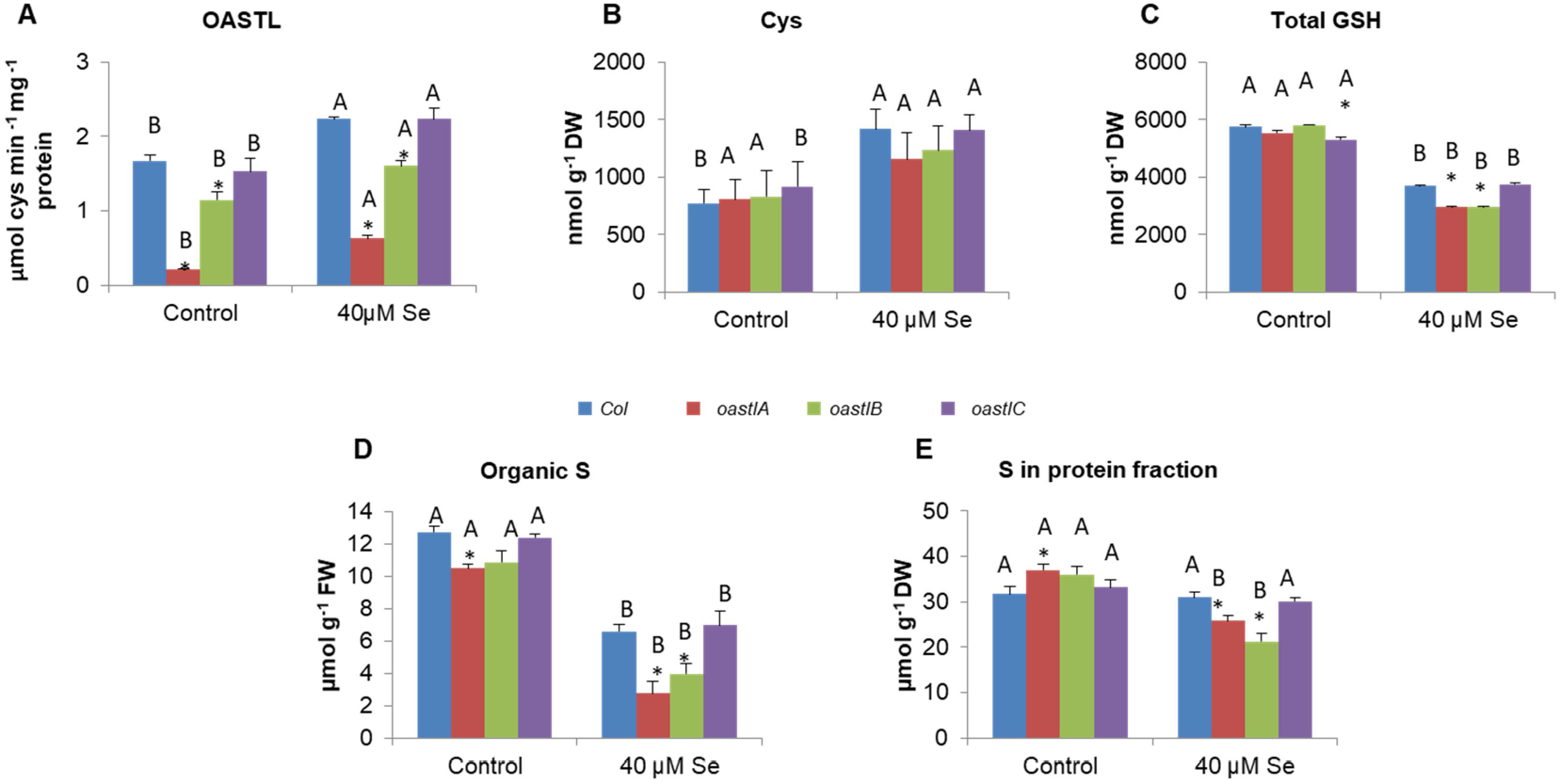
Cys biosynthesis and S containing compounds in WT, *oastlA, oastlB* and *oastlC* KO mutants supplemented with or without selenate for 14 days. The effect of 40 µM selenate treatment on OAS-TL activity (A), Cysteine level (B), Total glutathione (C), Total organic S (D) and S in protein fraction (E). The values are means ± SE (n = 4). Values denoted with different letters are significantly different according to T-test analyses, P < 0.05 (JMP 8.0 software, http://www.jmp.com/). Different uppercase letters indicate significant differences between control and Se treatment of each genotype. Asterisks indicate significant differences between WT and the KO plants *oastlA, oastlB and oastlC* subjected to the same treatment.

To examine this assumption the Cys, total glutathione and total organic S levels, as well as the S level in protein fraction were detected. No differences were noticed in cysteine level between WT and mutants in control and selenate treated plants, whereas in selenate treated plants a higher cysteine level was evident with WT and *oastlC* compared to these plants grown under control conditions (Fig. 2B). Determination of total glutathione revealed a decrease only in *oastlC* leaves whereas the other mutants exhibited comparable levels to WT leaves in plants grown in control conditions. In contrast, *oastlA* and *oastB* showed decreased total glutathione level compared to WT and *oastlC* leaves in plants exposed to selenate. Significantly, all genotypes exposed to selenate exhibited significantly lower glutathione levels than control plants (Fig. 2C). Determination of the organic S level revealed a significant reduction in all the genotypes at 40 µM selenate compared to untreated controls. Under the selenate stress, *oastlA* and *oastlB* exhibited a significant decrease in organic S as compared to WT and *oastlC* (Fig.2D).

The level of S in the protein fraction, under control conditions, was higher in *oastlA* than in WT, whereas in the other mutants it was similar to *WT*. Selenate supplementation resulted in a significantly lower S content in the protein fraction in *oastlA* and *oastlB* as compared to their controls and selenate treated WT and *oastlC* (Fig.2E). Taken together, the lower total glutathione, organic S and S in protein fraction in the leaf tissue of mutants impaired in OASTLA or B proteins fed with selenate (Fig. 2C, D and E), indicates possible interference in the S reduction pathway by selenate, resulting in retarded plant growth as compared to WT and *oastlC* mutant.

### The presence of selenate in the growth medium induces catabolic processes in *Arabidopsis*

Cysteine can be found in plants as free cysteine, being among the amino acids that build proteins, as well as an intermediate of Met biosynthesis, GSH and many other thiol species (Hesse and Hoefgen, 2003; Ravanel et al., 1998). Under the long day (14/10 light/dark) growth conditions free cysteine levels in leaves of plants growing in plates containing 0.5 MS medium, were similar in the mutants compared to WT either in control or in selenate treated plants (Fig. 2B). Considering the lowest cysteine generating activity level of OASTL was evident in *oastlA* and *B* (Fig. 2B), this result indicates a possible catabolic protein degrading activity which can act as an additional source of free L-cysteine in *oastlA* and *B*. In support of this notion is the significantly lower soluble protein content evident in *oastlA* and *B* compared to WT and *oastlC* mutant grown with selenate in the growth medium (Fig. 3A).

**Figure 3.**
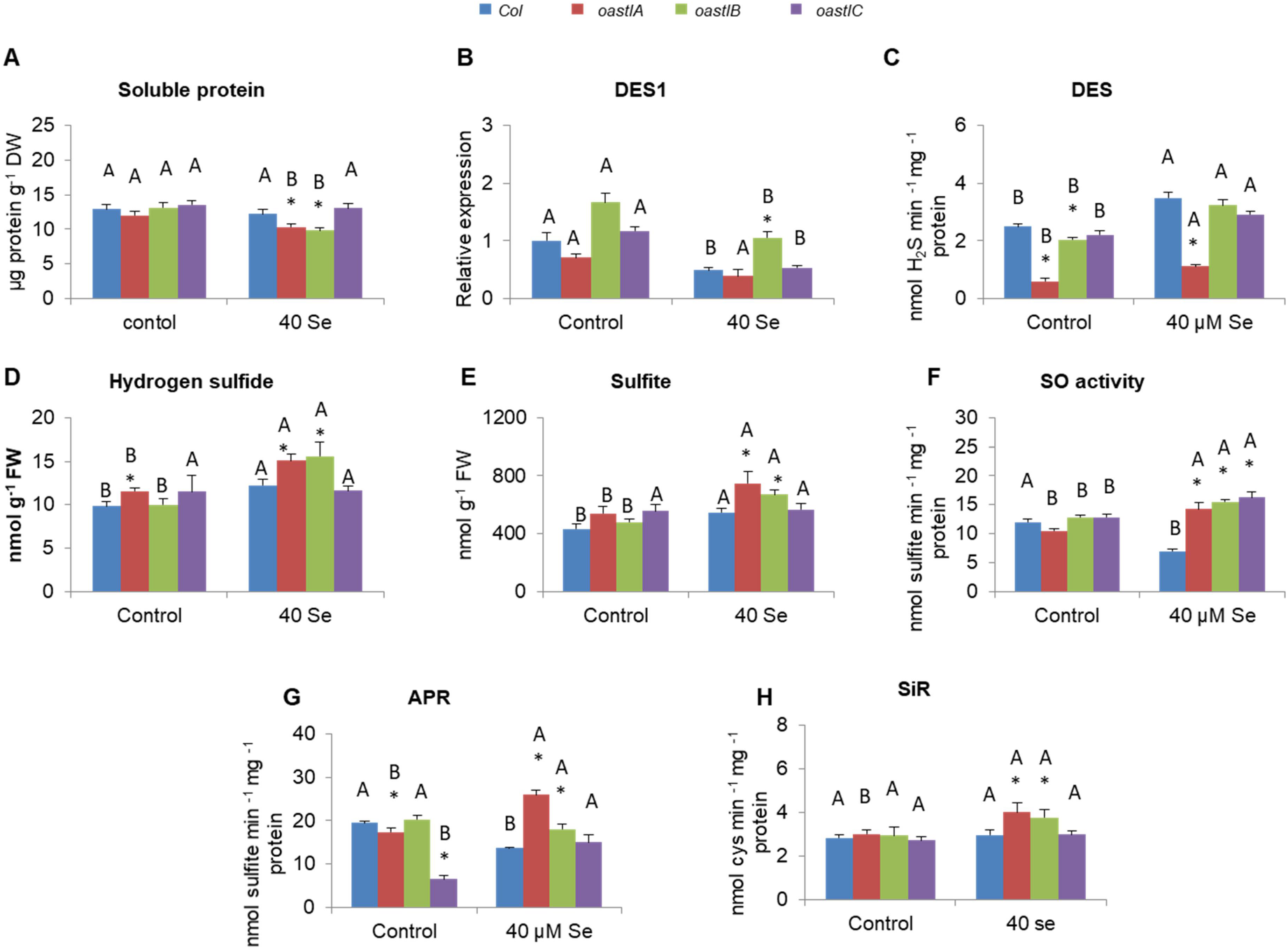
Soluble protein level and S metabolism components in WT, *oastlA, oastlB* and *oastlC* KO mutants supplemented with or without selenate for 14 days. The effect of 40 µM selenate treatment on soluble protein content (A), transcript expression of DES1 (B), L-Cysteine desulfhydrase (DES) activity (C), sulfide (D) and sulfite levels (E), Sulfite oxidase (SO) activity (F), Adenosine 5′-phosphosulfate reductase (APR) activity (G) and Sulfite reductase (SiR) activity (H). Transcript level of *DES1* was detected by quantitative reverse transcription-PCR using *ACTIN2* as the housekeeping transcript for normalization. The relative expression in the various genotypes was analyzed using the normalized WT control as reference. The values are means ± SE (n = 4). Values denoted with different letters are significantly different according to T-test analyses, P < 0.05 (JMP 8.0 software, http://www.jmp.com/). Different uppercase letters indicate significant differences between control and Se treatment of each genotype. Asterisks indicate significant differences between WT and *oastlA, oastlB, oastlC* plants subjected to the same treatment.

Recently, a lower biomass accumulation rate, the result of reduced organic S content was shown to be the consequence of higher L-cysteine degradation by desulfhydrase (DES) activity of OASTL in the perennial halophyte *Sarcocornia*, but not in the annual halophyte *Salicornia*, both fed with high sulfate (Kurmanbayeva et al., 2017). To examine whether OASTLs play a role in cysteine degradation also in non-halophyte plants, we studied first the expression of the cytosolic *DES1* claimed previously to play an important role in cytosolic L-Cys degradation (Álvarez et al., 2010). Unlike the *OASTLs* whose transcript levels were enhanced in WT plants in response to selenate (Supplemental Fig. S2 D-F), a significant decrease in *DES1* transcript abundance was evident in WT, *oastlB* and *C*, whereas in *oastlA* mutant a non-significant decrease was noticed. Under control and selenate stress conditions, *DES1* transcript abundance was the highest in *oastlB* (Fig. 3B).

Unlike the *DES1* transcript abundance in *WT* and *OASTLs* mutants, but similar to the L-Cys generation activities of the OASTLs (Fig. 2A), the total L-Cys DES kinetic activity was enhanced in all genotypes grown with selenate compared to unstressed control plants (Fig. 3C). DES activity rate was significantly lower in *oastlA* and *oastlB* mutants compared to *WT* grown under unstressed conditions, while when exposed to selenate *oastlA* exhibited a significant lower activity rate than *WT* and the other genotypes (Fig.3C). These results indicate a possible role of these OASTL isoforms in L-Cys degradation and not only in L-Cys biosynthesis (Figs.2A, 3C).

Notably, in spite of the lower or similar DES activity rate, *oastlA* and *oastlB* mutants exhibited higher sensitivity to selenate stress, accumulating lower biomass. This can be attributed to the lowest capacity of L-Cys biosynthesis by OASTL in *oastlA* and *oastlB* mutants grown with selenate that resulted in low levels of organic sulfur, total glutathione as well as low S in the protein fraction (Fig. 2A,C,D,E). This could result in S-starvation type responses such as the higher sulfide level evident in these mutants compared to WT and *oastlC* when stressed by selenate, exhibiting, excluding *oastlC*, higher sulfide than the control unstressed plants, (Fig. 3D). The higher sulfite level detected in these mutants compared to WT and *oastlC* grown with selenate (Fig. 3E), shown before when detected on fresh weight basis to be toxic in *Arabidopsis* and tomato plants (Brychkova et al., 2012; Brychkova et al., 2013; Yarmolinsky et al., 2014), is likely another S-starvation type response and an additional cause for the mutants’ sensitivity to selenate. Interestingly, while under control unstressed conditions SO activity was similar in mutants and in WT treated with selenate, the mutants exhibited a drastically higher SO activity rate than WT (Fig. 3F). This enhanced SO activity in the mutants compared to WT is a typical response of SO to sulfite increase as was shown before (Brychkova et al., 2012; 2013; Yarmolinsky et al., 2013; 2014). The selenate induced sulfite increase in *oastlA* and *oastlB* mutants is most likely the result of the increased APR activity rate in these mutants compared to WT and *oastlC* grown with selenate (Fig. 3G). The increase in APR activity rate in *oastlA* and *oastlB* leaves is an expected response to S-starvation (Lee et al., 2011 and references therein), as featured in the current study by lower glutathione, organic S and S in the protein fraction of these mutants compared to WT and *oastlC* (Fig. 2C,D,E). Further reinforcement for this notion is the significant increase in SiR activity resulting in the enhanced sulfide in *oastlA* and *oastlB* treated with selenate, whereas no differences between the genotypes were noticed in the unstressed plants (Fig. 3H, D). Taken together, the increase in SO, APR and SiR activity, followed by the enhanced sulfite and sulfide levels (Fig. 3D-H), indicates a futile anabolic S-starvation response to the selenate induced organic-S catabolism (Fig. 2C, D, E) in *oastlA* and *oastlB* compared to Wild-Type and *oastlC*.

### OAS-TLs degrades L-Cys and L-SeCys to H_2_S and H_2_Se respectively

The decrease in L-Cys desulfydrase kinetic activity in the *oastlA* and *oastlB* mutants compared to WT (Fig. 3C) raised the possibility that OASTLs may act as L-Cys and L-SeCys desulfhydrases. This was examined by employing crude protein extracted from leaves of 14 day old selenate stressed and unstressed WT, *oastlA, B* and *C* plants, fractionated by native SDS-PAGE (Sagi and Fluhr, 2001; Srivastava et al., 2017; Yesbergenova et al., 2005) to distinguish between the activity bands of the various OASTLs. The gels were then subjected to DES activity reaction solution containing L-Cys or L-SeCys as a substrate under reducing conditions, generated in the presence of β-Mercaptoethanol. The reaction of lead (as lead acetate) with sulfide or selenide (the consequence of desulfhydration activity that degrades L-Cys or L-SeCys, respectively), results in a brown-dark precipitate exhibiting the activity band at the position of the fractionated active enzyme. When the substrates (L-Cys or L-SeCys) or reducing conditions (β-Mercaptoethanol) were omitted from the reaction solution, the activity bands, the precipitated product of the generated lead sulfide or lead selenide, were not seen (Supplemental Fig. S3A). In contrast, when the substrate was present under reducing condition, three distinguishable activity bands were present in the location of the fractionated WT proteins (Fig.4A). Significantly, the fractionated proteins extracted from *oastlA, oastlB* and *oastlC* mutants exhibited the absence of the lowest, middle and most upper activity band respectively, compared to WT, with L-Cys as the substrate (Fig.4A). When L-SeCys was present in the reaction solution as the substrate, the two lowest activity bands with the same mobility as with the L-Cys desulfhydrase activity were present in the fractionated WT proteins, whereas the fractionated proteins of *oastlA* and *oastlB* mutants were missing the lowest and middle bands, respectively (Fig. 4B). The absence of the most upper activity band of OASTLC in WT and *oastlA, oastlB* mutants, is likely the result of the relatively low L-SeCys desulfhydrase activity rate by OASTLC, being below the detection level when employing L-SeCys as a substrate. Importantly, the presence of selenate in the growth medium resulted in the general enhancement of L-Cys and L-SeCys desulfhydrase activities of OAS-TL enzymes compared to the control conditions (Fig.4A and B). These results indicate that OASTLA, OASTLB and OASTLC play a role not only in Cys/SeCys biosynthesis, but also in degrading L-Cys and L-SeCys.

**Figure 4.**
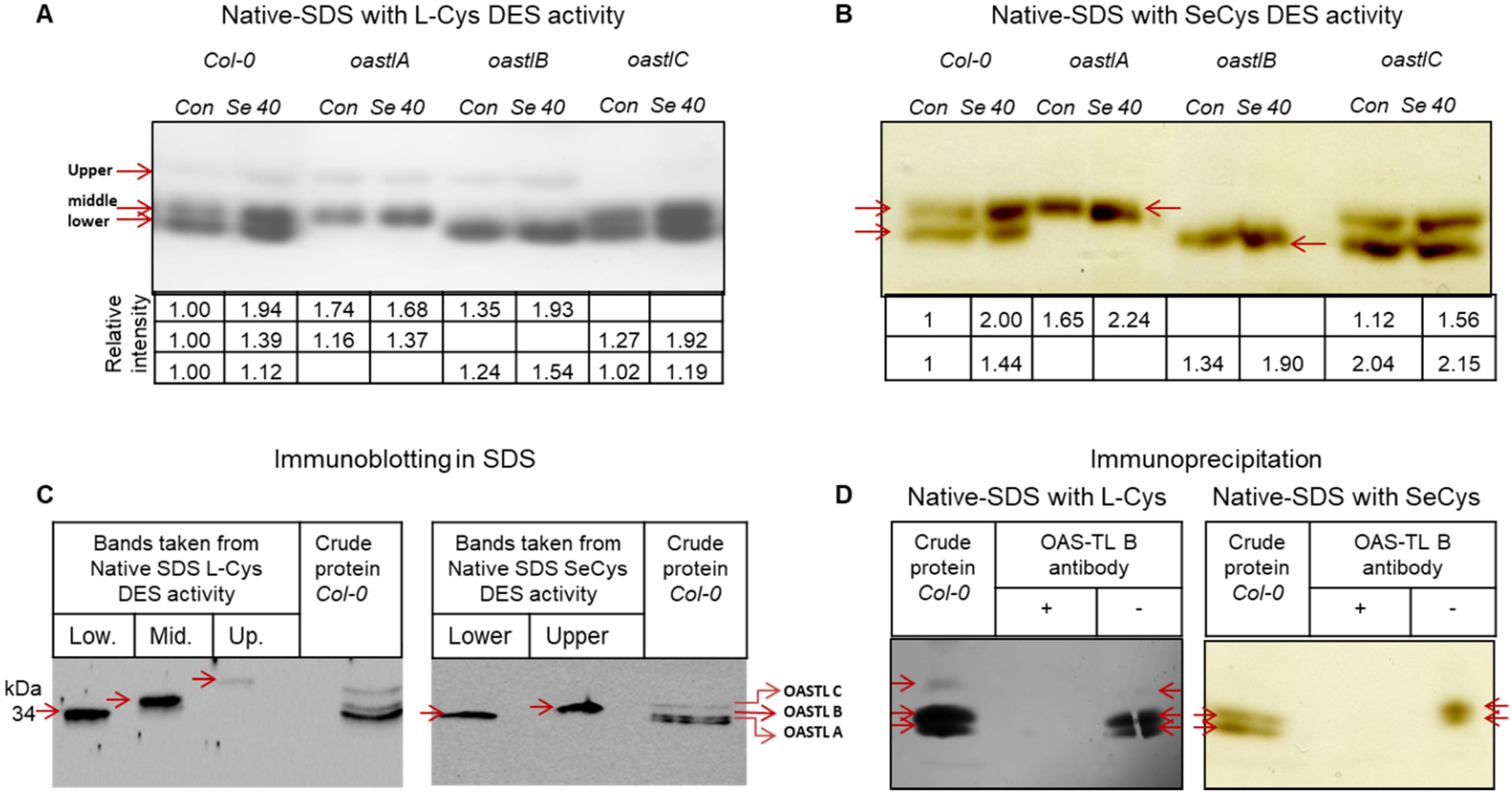
NATIVE-SDS on L-Cys and SeCys desulfhydrase activities in WT, oastlA, oastlB and oastlC KO mutants supplemented with or without selenate for 14 days. Effect of 40 µM selenate treatment on L-Cysteine (A) and SeCys (B) degradation activities. C. Immunodetection of the sliced WT activity bands from 4A are in left insert and the sliced WT activity bands sliced from 4B are in the right insert. D. Desulfhydrase activity of WT crude protein extract without immunoprecipitation (left lane) with immunoprecipitation in the presence (+) or absence (-) of specific antibodies. L-cys and L-Secys desulfhydrase activities are left and right inserts respectively. Red arrows indicate activity band.

Interestingly, while the homozygous *Atdes1* KO T-DNA mutant SALK_103855 was shown to express ca 45 to 60% of *DES1* (AT5G28030) compared to WT root transcript (see Figure 7C in Mei et al., 2019), the homozygous KO mutant SALK_103855 exhibited almost the absence and ca 29% transcript expression compared to WT employing oligonucleotides upstream and downstream along the transcript length, respectively. SALK_205358, shown before to be a *des1* null mutant (Jin et al., 2018) exhibited ca 40% decrease in *DES1* compared to the expression in WT leaves, employing the oligonucleotides described above (Supplemental Fig. S3B and E). Yet, in-spite of the transcript decrease the *Atdes1* KO (SALK_205358) exhibited kinetic activities of cysteine generation by OASTLs and total desulfhydration activity in control and selenate stressed plants comparable to WT (Fig. S3C and D, respectively).

Since SALK_103855, KO mutant of *Atdes1* grown on MS medium was also demonstrated to be responsible for only a minor portion of the Cys-desulfudrase activity compared to WT [14% of the total activity in leaves of plants grown in plates containing agar with MS medium (Alvarez et al., 2010)], an in-gel desulfydrase activity was employed to further examine the significance of DES1 and the OASTLs in desulfhydrase activity in Arabidopsis plants. NATIVE-SDS was employed to fractionate the proteins (Sagi and Fluhr, 2001; Yesbergenova et al., 2005) for desulfhydrase in-gel activities in WT and homozygous SALK_205358 *Atdes1* mutant employing L-Cys or L-SeCys as substrates. The existence of OASTLA, OASTLB and OASTLC activity bands with relative intensity in *Atdes1* mutant as with WT was evident (Supplemental Fig. S3E). We further examined L-Cys desulfhydrase activity in SALK_103855 leaves and further noticed similar activity bands in WT and the *Atdes1* homozygous mutant (Supplemental Fig. S3F, left insert). These results demonstrate that the OASTL enzymes may play a significant role in degrading L-cys and L-SeCys to sulfide and selenide respectively in Arabidopsis leaves.

To further support the identification of the enzymes which generated the activity bands, a Western blot analysis employing the lower middle and upper sliced activity bands in WT (Fig. 4A and B) was performed after the fractionation of the proteins in SDS-PAGE, using OASTL C antibody which cross reacts also with OAS-TL A and B proteins (kindly provided by Prof. R. Hell, Heidelberg University, Germany). Different mobility of the single bands was demonstrated (Fig.4C left side), supporting the notion of OASTL A, B and C in L-Cys and OASTL A, B isoforms playing a role in L-SeCys degradation (Fig.4C right side). Additional support for this notion was achieved by immunoprecipitation analysis. The proteins that generated L-Cys and SeCys DES activity bands were fully pulled down after the proteins were immunoprecipitated by *Arabidopsis* OAS-TL B specific antibody (kindly provided by Prof. Dr. S. Kopriva, University of Cologne, Germany) that also cross reacts with OASTLA and C isoforms (Fig. 4D left and right side respectively).

Finally, the identification of tripsinized unique peptides in the three sliced activity bands after the fractionation of WT crude protein extract (Fig.4A and B) was performed. The lowest sliced activity bands of L-Cys and L-SeCys desulfhydrase activities revealed 23 and 20 unique peptides overlapping 66% and 65% of the Arabidopsis OAS-TL A protein sequence, respectively (Supplemental Table S1A-C). Together with the OAS-TL A peptides, a lower amount of 14 and 9 peptides of OASTLB were identified in the lowest sliced L-Cys and SeCys desulfhydrase activity bands, respectively (Supplemental Table S1A-C). In the middle activity band, sliced from L-Cys and SeCys desulfhydrase activity, 23 and 18 unique peptides were identified showing 58% and 60.5% overlap of OAS-TL B protein sequence, respectively (Supplemental Table S1A-C). Additionally, 18 and 15 peptides of OASTLA proteins in the L-Cys and SeCys desulfhydrase activity bands and 6 unique peptides of OASTLC protein were identified in L-Cys desulfhydrase activity. The upper band sliced from L-Cys desulfhydrase activity revealed 13 specific unique peptides with 45% coverage of OASTLC protein. Six and 11 unique peptides of OASTLA and B proteins respectively were identified as well (Supplemental Table S1A-C). The lack of a full separation between the 3 OASTLs activity bands is likely the result of the relatively high identity between the three proteins [ca 70 to 80% identity (Supplemental Fig.S4)] that prevented the clear complete separation in-spite of the 16 cm fractionation length by SDS-PAGE. Yet, based on the identification by the use of the three mutants exposed to in-gel activity, immunoblot and immunoprecipitation by the specific antibodies employed (Fig. 4), as well as the majority of the overlapping unique peptides (Supplemental Table S1), we can conclude that OASTLA, B and C are localized to the highest, second-highest and the lowest activity bands, respectively of L-Cys and SeCys desulfhydrase activity.

Significantly, only two unique peptides of DES1 were identified in the lower and middle bands and none in the most upper band of L-Cys desulfhydrase activity bands (Supplemental Table S1). Considering that the number of theoretic tryptic peptides in DES1 is similar to that of OASTLs (estimated by the use of Expasy: https://web.expasy.org/peptidemass/), these results indicate that DES1 is a less abundant protein or it undergoes ionization. Yet, since the semi quantitation was made by calculating the peak area of each peptide, whereas to overcome differential ionization, the area of the protein was calculated as the average of the three most intense peptides from each protein; the data indicate that OASTL A, B and C are more abundant than DES1 protein (Supplemental Table S1D).

The less abundant DES1 protein in the activity bands does not necessarily indicate decreased L-Cys and L-SeCys DES activity by the WT DES1 compared to the OASTLs. Yet, firstly, the kinetic activity revealed a major significant decrease of 80 and 65% in the Cys-desulfhydrase activity of the cytosolic *oastlA* leaves in plants grown in control or selenate stressed plants, respectively compared to WT (Fig.3C). Additionally, activity bands, additive to OASTL, A, B and C, that could be attributed to DES1, were not noticed, neither above, nor below and none were noticed at the position of the migrated OASTL A, B and C when the activity bands were absent in the related impaired mutants under control and selenate stressed conditions (Fig. 4A and B). Taken together these results (Supplemental Table S1A-D, Figs 3C, 4A, B) strongly support the notion that OAS-TL A, B and C proteins play a major role in L-Cys and L-SeCys degradation in *Arabidopsis*. In the presence of selenate, plants impaired in OASTL A and B exhibit higher catabolic processing rates of chlorophyll and protein degradation, the result of enhanced sulfur metabolism interference in the mutants compared with WT.

## DISCUSSION

### Impairment in OASTLA and OASTLB leads to higher sensitivity to selenate

Non-specific incorporation of seleno-amino acids SeCys and SeMet into proteins instead of cysteine and methionine is thought to be a main cause for Se induced toxicity in many plants (Sors et al., 2005; Terry et al., 2000). Yet, lower Se in protein fraction was evident in *apr2* KO mutants exhibiting stronger sensitivity than WT to the Se treatment (Grant et al., 2011). Similarly, determination of the total Se and Se content in protein fraction revealed a significantly lower rate in *oastlA* and *oastlB* mutants, compared with WT and *oastlC* (Fig. 1E and Supplemental Fig. S2B). This indicates that the accumulated Se in *oastlA* and *oastlB* mutant proteins is not the main cause for their stronger sensitivity to Se compared with WT.

The uptake and assimilation of Se has similar effects on gene expression as with the case of S-starvation, resulting in greater S uptake and assimilation rates by plants [(White, 2018) and references therein]. Plants exposed to high Se level exhibited symptoms such as growth inhibition and chlorosis (Grant et al., 2011; Sabbagh and Van Hoewyk, 2012; Van Hoewyk, 2018). Interestingly, the overexpression of OASTL in Arabidopsis and tobacco increased tolerance to the toxicity of Cd stress by increasing thiol availability (Domínguez-Solís et al., 2004). Among the three *Arabidopsis* OASTL mutants grown for 2 weeks with 40µM selenate, a decrease in biomass accumulation and chlorophyll content compared to WT was noticed in *oastlA* and *oastlB* but not in *oastlC* mutant (Fig.1). This is likely since the cytosolic and chloroplastic OASTL A and B respectively, exhibit significantly higher activity rates compared with the mitochondrion localized OASTL C, the latter being responsible for only 5-12% of OASTL activity in WT plants (Heeg et al., 2008; Kuske et al., 1996). The results demonstrate that dissimilar to WT and impairment in OASTLC, in the absence of active OASTLA and OASTLB, the selenate interference in S assimilation pathway leads to a stronger decrease in organic S, S in protein fraction and total glutathione, and has a more negative effect on the remaining chlorophyll level and the resulting biomass accumulation than the expected positive effect of lower selenium content in the organic fraction (Figs 1 and 2).

The free cysteine levels were similar in WT and the mutants exposed to selenate (Fig.2B), whereas the Cys generation activity rate differed as shown here in response to selenate (Fig.2A). This was also shown previously in an O-acetylserine (thiol) lyase reduced isoform by RNA interference in potato, as well as in *Arabidopsis* KO mutants (Heeg et al., 2008; Riemenschneider et al., 2005), indicating the feasibility of an additional Cys source affecting free Cys level. The source can be protein degradation, indicated by the lower soluble protein level in *oastlA* and *oastlB* KO mutants compared to WT and *oastlC* treated with selenate (Fig. 3A). The protein degradation is the result of selenate induced stress, being stronger in *oastlA* and *B* as indicated by the lower remaining chlorophyll (Fig. 1D), organic-S, and S in protein fraction (Fig. 2D and E).

Sulfite is a strong nucleophile that should be tightly regulated to avoid its deleterious reaction with a wide variety of cellular components (Brychkova et al., 2007; Lang et al., 2007; Brychkova et al., 2013; Yarmolinsky et al., 2013; Brychkova et al., 2015). The increased sulfite level in *oastlA* and *oastlB* mutants compared to WT (Fig. 3D) is a deleterious effect of a futile S-starvation response, the result of the low organic sulfur, total glutathione as well as low S in the protein fraction (Fig. 2A,C,D,E) in *oastlA* and *oastlB* mutants treated with selenate. APR activity increase in *oastlA* and *oastlB* leaves that caused the enhanced sulfite level in these 2 mutants is a response to S-starvation (Lee et al., 2011) and the followed sulfide level increase, the result of higher SiR activity in these 2 mutants, further supports the notion of selenate induced S-starvation. The futileness of such response is firstly because of the waste of reducing agents and ATP (see in Supplementary Fig. S1) in the process ending with excess sulfide, the result of the inability of *oastlA* and *oastlB* plants to assimilate sulfide to L-Cys generation. Additionally, the sulfite level increase, shown before to induce oxidative stress (Brychkova et al., 2012; Yarmolinsky et al., 2014), indeed led to oxidative stress (Supplementary Fig. S5) that had an additive deleterious effect on the growth of *oastlA* and *oastlB* mutant plants. The oxidative stress could further induce APR to generate more sulfite as demonstrated before (Bick et al., 2001; Koprivova et al., 2008). Notably, the levels of hydrogen peroxide (H_2_O_2_), the carbonyl toxic malondialdehyde (MDA), the product of lipid peroxidation by prolonged oxidative stress (Tian et al., 2017) and the antioxidant anthocyanins (Gould et al., 2002; Xu et al., 2017), were enhanced (on FW, and except H_2_O_2,_ on soluble protein basis as well) in the selenate treated plants compared to control plants. MDA and anthocyanin levels were significantly higher in *oastlA* and *oastlB* compared to the selenate treated WT and *oastlC* mutant, whereas, H_2_O_2_ was significantly higher in the 3 mutants compared to WT treated with selenate. The enhancement in H_2_O_2,_ MDA and anthocyanins levels in *oastlA* and *oastlB* mutants (Supplemental Fig. S5) is indicative of the higher oxidative stress level in these plants as shown before (Gould et al., 2002; Srivastava et al., 2017; Xu et al., 2017; Yarmolinsky et al., 2014). Taken together the results indicate that the decrease in *oastlA* and *oastlB* biomass accumulation is likely the result of selenate induced S-starvation responses causing waste of reducing agents and ATP, as well as sulfite induced oxidative stress.

### OASTL A and B act as major L-Cys and L-SeCys desulfhydrases in *Arabidopsis*

The similar free cysteine level in WT and *oastlA* and *B* mutants, compared with the lower cysteine generating activity rate of OASTL and the lower soluble protein level evident in the mutants exposed to selenate (Figs 2A, B, Supplemental S3A), led us to examine a role for OASTL proteins in degrading L-Cys and SeCys to sulfide and selenide respectively. Additionally, albeit in a small rate comparing to the biosynthesis by OASTLs, the degrading activity of the active enzymes is likely beneficial to plants, decreasing the levels of L-SeCys (compare the rate in Fig. 2A to the rate in Fig.3C) and its incorporation into proteins.

L-Cys desulfhydrase activity in WT showed 3 activity bands, where the upper band intensity was weaker than that of the other two bands. L-SeCys desulfhydrase exhibited only 2 activity bands, with the absence of the upper band likely the result of detection limits (Fig. 4A and B). Importantly, fractionated proteins extracted from *oastlA* and *oastlB* KO mutants exhibited in-gel activity absent of the lower and middle activity band, respectively with both activity types; the L-Cys and SeCys DES, whereas the most upper band was seen only when L-Cys was used as the substrate and only in WT, *oastlA* and *oastlB* but not in *oastlC* (Fig. 4A and B). These results indicate that the activity bands with the highest, second-highest and the lowest mobility are of OASTLA, B and C, respectively. Additionally, the results show that the three OASTLs have the capacity of L-SeCys and L-Cys DES activities. This was further supported by the use of specific antibodies to identify the sliced activity bands by Western blot after further SDS fractionation (Fig. 4C), as well as by immunoprecipitation assay (Fig. 4D). Finally it was supported by the identification of the tripsinized unique peptides (Supplemental Table S1) in the three sliced WT activity bands (Fig.4A and B).

Importantly, DES1 was shown to degrade L-Cys, estimated by the use of the *des1* null mutants to be responsible for up to 14% of the total L-Cys desulfhydrase activity compared to WT plants grown in plated on MS medium (Álvarez et al., 2010). This suggests the existence of additional source/s that can contribute to the majority (ca 86%) of H_2_S production by degrading L-Cys. Notably, the impairment of *oastlA* and *oastlB* mutants led to a significantly decreased sulfide production activity rate by L-Cys desulfhydrase [ca 76% and 18% reduction respectively (Fig3C)] compared to WT, indicating the important role of the OASTLs in degrading L-Cys. Intriguingly, only very few unique peptides of DES1 were identified in the major activity bands, the highest and second-highest mobility bands, of L-Cys desulfhydrase activity and none in the SeCys desulfhydrase activity bands, indicating that the OASTLs are more abundant proteins than DES1 (Supplemental Table S1). Taking together i) the lower abundancy of DES1 compared to the OASTLs and ii) the significant decrease in the Cys-desulfhydrase kinetic activity of the *oastlA* leaves in plants grown in control or selenate stressed plants compared to WT (Fig.3C) as well as iii) the absence of activity bands, additive to OASTL A, B and C activity bands that could be attributed to DES1 in control and selenate stressed condition (Fig. 4A and B), the result strongly support the notion that OASTL A, B and C play a major role in L-Cys and L-SeCys degradation in *Arabidopsis*.

## CONCLUSIONS

Using *Arabidopsis* O-acetylserine-(thiol) lyases mutants, impaired in the biosynthesis of cysteine (Cys) from the substrates sulfide and O-acetylserine, it was shown that the absence of active OASTLA and OASTLB, confers higher reduced growth, lower remaining chlorophyll, lower sulfur in protein fraction and lower total glutathione than in selenate stressed WT and *oastlC* mutant as the result of the interference of selenate with the sulfate reduction pathway. Additionally it was shown that the absence of active OASTLA or OASTLB confers higher sensitivity to selenate compared to WT or the absence of active OASTLC. Further, it was shown here that the cytosolic OASTLA, chloroplat localized OASTLB and the mitochondrion localized OASTLC has an important role in desulfhydrase activity degrading not only L-Cys but also SeCys to sulfide and selenide, respectively.

## MATERIALS AND METHODS

### Plant Material, Growth Conditions and Selenium treatment

*Arabidopsis thaliana* var. Columbia wild type (*WT*), *oastlA (Salk_072213), oastlB (Salk* _*021183), oastlC (Salk_000860) and des1* (SALK_205358) lines were used for Selenium treatment. SALK_103855 (*des1*) was employed for in-gel DES activity and transcript expression. All the mutants were derived from the Columbia (*Col*) ecotype. Experiments were carried out in the growth room at Sde Boqer Campus, Ben-Gurion University of the Negev, under 14 h light/10 h darkness, 22°C and 75–85% relative humidity under photosynthetically active radiation of 150 µmol m^-2^ sec^-1^ as described in Brychkova et al., 2007.

Seeds were germinated and grown in standard 90 mm Petri dishes on solid 1/2 MS medium for 9 days (2 days in dark and 7 days in growth room). The seedlings were transferred to large Petri dishes of 155×30 mm diameter and height respectively, supplied with 1/2 MS medium (Murashige and Skoog, 1962) supplemented with or without 40µM of Na_2_SeO_4_. All treatments were performed in six replicates. The weight of shoot biomass accumulation was determined 14 days after the treatment and results were expressed as average plant growth rate in mg plant^-15^

### RNA Isolation, cDNA Preparation, and transcript analyses by Real-time PCR

To quantify the transcripts using quantitative reverse transcriptase-polymerase chain reaction (qRT-PCR), total RNA was prepared using the Aurum^TM^ total RNA Mini Kit (Bio-Rad, Hercules, CA) according to the manufacturer’s instructions. The cDNA was prepared in a 10 μl volume containing 350 ng of plant total RNA that was reverse-transcribed with an iScript^TM^ cDNA Synthesis Kit using modified MMLV-derived reverse transcriptase (Bio-Rad), a blend of oligo-d(T) and random hexamer primers, according to the manufacturer’s instructions. The generated cDNA was diluted 10 times and the quantitative analysis of transcripts was performed employing the sets of primers from both sites of insertions shown in Supplemental Table S2 as previously described (Brychkova et al., 2007). The transcripts level detected by Real-time PCR were normalized using *ACTIN2* (At3g18780) and *ELONGATION FACTOR1-α* (At5g60390) and compared to WT grown under the control condition. The data are presented as relative expression (means ± SE, *n* = 4). All PCR fragments were sequenced for verification.

### Chlorophyll, MDA, H_2_O_2_ and Anthocyanin determination

For detection of remaining content of chlorophyll (%) the leaf discs (7 mm diameter) were immersed in 90% ethanol and incubated at 4°C for 2 d in the dark. Absorbance of the extracted chlorophyll was measured at 652 nm, and total chlorophyll was estimated (Ritchie, 2006). MDA levels were determined as described by Srivastava et.al., 2017.

For detecting H_2_O_2_, frozen leaves were extracted in 50 mM P buffer (pH 7.5) at a ratio of 1:8 (w/v) and centrifuged (Eppendorf 5417R) twice at 18,000*g* for 20 min. The reaction mixture for detecting H_2_O_2_ consisted of 0.85 mm 4-aminoantipyrine, 3.4 mm 3,5-dichloro-2-hydroxobenzene sulfonate, 4.5 U mL^−1^ HRP in 2 mL of 50 mM P buffer (pH 7.5) in the presence or absence of 2 mM tungstic acid and 100 μM DPI as described in Yesbergenova et al. 2005. Absorbance was measured after 5 min at 515 nm as described above.

The anthocyanin content was determined based on a modification of protocols described by Laby et al. (2000) and Kant et al. (2006). Approximately 100 mg of fresh plants grown in 1/2 MS medium were crushed in 600 µL methanol acidified with 1% HCl. The extract was centrifuged for 10 sec at 4000g. Five hundred µL of double distilled water were added to the collected sand, mixture was gently vortexed and then 700 µL chloroform was added and mixed for 20 sec followed by centrifugation at 4000 g for 2 min. The total anthocyanin in the aqueous phase was determined by detecting the optical density (OD) at A530 and A657 nm. The amount of anthocyanin was calculated by subtracting the A657 from the A530 (Laby et al., 2000).

### Enzyme activity measurements

Protein extraction, desalting, concentration determination and kinetic assays for APR, SO, SiR, OAS-TL and DES activities were assayed and expressed as described before; OAS-TL and SiR activity in nmol Cys mg^−1^ protein min^−1^, APR and SO in nmol sulfite mg^−1^ protein min^−1^ and DES activity in nmol H_2_S mg^−1^ protein min^−1^ (Brychkova et al., 2012a; Brychkova et al., 2012b; Kurmanbayeva et.al., 2017). In brief, APR activity employing APS as substrate was detected using the sulfite-specific fuchsine colorimetric detection method. SO activity was measured as the disappearance of sulfite. The desalted protein extracts were treated with 1 mM tungstate for 30 min at 4°C to inhibit SO activity. SiR activity was estimated by the coupled SiR/OASTL assay with the addition of NADPH and tungstic acid. The resultant generated Cys was detected as described before (Gaitonde, 1967; Brychkova et al., 2012b). OAS-TL activity was initiated by incubating 75 μL Reaction mixture (RM) containing 25 mM OAS, 1 mM DTT, 1.42 mM Pyridoxal-phosphate (PLP), 60 mM Na_2_S and 200 mM P-buffer (pH 7.5) with 25 μL of protein extract for 10 min at 30°C. Generated Cysteine was detected at 560 nm by a spectrophotometer (Gaitonde, 1967).

DES activity was detected as H_2_S formation in the presence of L-cysteine. The assay solution contained 0.1 M Tris-HCl, pH 9.0, 2.5 mM dithiothreitol, 0.8 mM L-cysteine and 10 µg of desalted protein in a total volume of 0.2 mL. After incubation at 37 °C for 30 min, H_2_S was detected according to Bloem et al., 2004 with 30 mM FeCl_3_ dissolved in 1.2 N HCl and 20 mM N,N-dimethyl-p-phenylenediamine dihydrochloride dissolved in 7.2 N HCl. The formation of methylene blue was measured at 670 nm. Na_2_S*9H_2_O was used as a standard.

### Protein extraction, fractionation for in gel DES activity

Whole protein from Arabidopsis seedlings was extracted as described by Sagi and Fluhr, 2001. Concentrations of total soluble protein in the resulting supernatant were determined according to Bradford, 1976. Native-SDS PAGE was carried out as follows: samples were subjected to a Bio-Rad Protean II xi slab cell (Bio-Rad, Richmond, CA, USA), with a discontinuous buffer system (Laemmli, 1970) in 7.5% (w/v) polyacrylamide separating gels and 4% (w/v) stacking gels. Native-SDS PAGE was carried out using 1.5 mm thick slabs loaded with 200 µg of proteins. Regeneration of the active proteins after native-SDS PAGE was carried out by removal of the SDS as described by Sagi and Fluhr, 2001 with slight modification: by shaking the gel for 1 h in 10 mM Tris-HCl buffer (pH 7.8) solution (65 ml buffer per ml of gel) containing 2 mM EDTA.

DES activity was detected using a modification of an in-gel visualization protocol for H_2_S (Manchenko, 2002). Lead acetate was employed to detect the generated H_2_S and possibly H_2_Se, producing dark brown lead sulfide/selenide bands. The reaction solution contained 0.15 M Tris–HCl, pH 8.5, 1 mM Dithiothreitol (DTT), 50 mM β-mercaptoethanol, 5 mM L-cysteine or 2.5 mM SelenoCystine, 0.1 mM pyridoxal 5-phosphate (PLP), 0.4 mM lead acetate. The reaction was stopped by immersion of the gel in double-distilled water.

### Western Blot and Immunoprecipitation Analysis

Western blot proteins were separated by SDS–PAGE carried out in 12.5% polyacrylamide gels and were blotted to polyvinylidene difluoride membranes (Immun-Blot membranes, Bio-Rad). The blotted proteins were subjected to immuno-detection with antibodies raised against Arabidopsis chloroplastic OAS-TL C (diluted1:1000) (kindly provided by Prof. R.Hell, University of Heidelberg, Heidelberg) and followed by secondary antibodies as described by Heeg et al., 2008.

For the immunoprecipitation assay 200 µg of protein from *WT* were incubated with 80 µl of the OAS-TL B (diluted1:5000) (kindly provided by Prof. Dr. S. Kopriva, University of Cologne, Köln) antibodies in Tris-buffer saline (TBS) for 30 min at room temperature and then kept at 4°C for overnight. Protein extract that had not been mixed with antibody was employed as control. The mixture and the control solutions were incubated with 120 µl of Protein G Agarose at 4°C for 2h with continuous shaking and then centrifuged at 10,000g for 5 min, followed by removal of supernatant for analysis by the Native-SDS DES assay.

### Determination of selenium and sulfur-containing metabolites

To determine the total selenium content, 100 mg of the dried and powdered leaves were placed in glass tubes, digested with 70% HN0_3_, heated at 220°C for 4 hours and quantified by inductively coupled plasma emission spectrometry (ICP-AES) (Kalra, 1997).

Se in protein fraction: 400 g fresh weight was extracted in 2 mL buffer containing 100 mM NaCl, 50 mM Tris HCl (pH 7.5), 0.5% (v/v) Triton X-100, 1 mM DTT, and 0.1 mM PMSF. The homogenate was cleared by centrifugation (7,500g for 10 min). A small sample was taken for protein determination, and the volume of the extract was measured. The proteins in the extract were precipitated by adding TCA to a final concentration of 15% (w/v). The mixture was incubated on ice for 30 min and then centrifuged for 20 min at 7,000g at 4°C. The pellet was washed with ice-cold acetone, dried, and dissolved in 1 mL of concentrated nitric acid. After acid digestion, the Se was determined by ICP (Pilon-Smits et al., 1999).

Selenate detection was done according to (Schiavon et al., 2012) with slight modifications. The frozen tissues (200 mg) were ground in liquid nitrogen, and then 5 mL of distilled water was added. The obtained extracts were filtered (0.22μm, Millipore) and analyzed for sulfate and selenate concentrations by Ion chromatography using a Dionex IonPac AS19 4×250mm column.

Organic S and Se were calculated by subtracting the sum of sulfate/selenate, sulfite and sulfide from the total sulfur/selenium content.

Determination of sulfite, free Cys and total glutathione was performed as described (Brychkova et al., 2013; Yarmolinsky et al., 2014). Sulfide was extracted (1:4, w/v) in 0.1 M Tris–HCl, pH 8.0 buffer in the presence of 0.5 M sodium sulfate to minimize sulfite self-oxidation to sulfate. After centrifugation at 1800 g for 15 min, the supernatant was further de-proteinized employing Sephadex G-25 Column (Pd-10, GE Healthcare) loaded with the same extraction solution. Immediately after the separation through the column 600 µl of 2% cadmium acetate were added to the 600 µl of the sample and kept on ice. The levels of sulfide were detected by 40 µl NN-Dimethyl-1,4-phenylen-diammonium-dichloride dissolved in 7.2 N HCl and 4 µl FeCl3 dissolved in 1.2 N HCl from 240 µl of preincubated sample with 2% cadmium acetate (Siegel, 1965). Sulfide was detected at 625 nm.

### Protein sequencing

To identify the proteins participating in the cysteine desulfhydrase activity, the activity bands from the in-gel activity of DES were sliced from the native gel, and fractionated with 12.5% SDS-PAGE (Fig. 4). Thereafter the proteins were stained by Coomassie Brilliant Blue, and the stained bands were excised from the gel, trypsinized and the resulting peptides were separated by HPLC and analyzed by LC-MS/MS on Q-Exactive (Thermo) at The Smoler Protein Research Center (Technion University, Haifa, Israel).

All the identified peptides were filtered with high confidence, top rank, mass accuracy, and a minimum of 2 peptides. High confidence peptides passed the 1% FDR threshold. (*FDR =false discovery rate, is the estimated fraction of false positives in a list of peptides). Semi quantitation was done by calculating the peak area of each peptide. The area of the protein is the average of the three most intense peptides from each protein. Analysis of peptide sequences was performed by employing Proteome Discoverer(tm) Software ver. 1.4.1.14 (Thermo Fisher Scientific Inc. https://www.thermofisher.com/order/catalog/product/IQLAAEGABSFAKJMAUH).

### Accession Numbers

Sequence data for this article can be found in the Arabidopsis Genome Initiative or GenBank/EMBL databases under the following accession numbers: At4g14880 *(OASTL A*), At2g43750 (*OASTL B*), At3g59760 (*OASTL C*), AT5G28030 (*DES1*), AT3G18780 (*ACTIN 2*), AT5G60390 (*EF 1-α*)

## ACKNOWLEDGMENT

We thank Prof. Dr. S. Kopriva (University of Cologne, Köln) and Prof. R.Hell, (Universität Heidelberg, Heidelberg) for providing OAS-TL A and B antibodies.

## Supplemental Figures

**Supplemental Figure S1**. Schematic model of Sulfur (S) and Selenium (Se) metabolism in Arabidopsis plants [after White (2015)]. ATP sufurylase (ATPS) catalyzes the adenylation of sulfate/selenate to adenosine 5′-phosphosulfate (APS)/adenosine phosphoselenate (APSe) using ATP as an electron donor. Then, APS/APSe is reduced by the plastidic enzyme APS reductase (APR) to sulfite/selenite in the presence of two molecules of reduced glutathione, which acts as an electron donor. The generated sulfite/selenite can be reduced to sulfide/selenide by the Sulfite Reductase (SiR) employing 3 molecules of reduced ferredoxin. Alternatively, selenite can also be reduced non-enzymatically by glutathione (GSH) to selenide. The sulfide/selenide together with O-acetyl-L-Serine (OAS) are the substrates for Cysteine (Cys)/Selenocysteine (SeCys) biosynthesis catalyzed by O-acetylserine-(thiol) lyase (OAS-TL).

**Supplemental Figure S2**. OASTLs protein and transcripts expression, as well as Se containing compounds in *WT, AtoastlA, AtoastlB* and *AtoastlC* KO mutants supplemented with or without selenate for 14 days. Effect of 40µM Selenate treatment on OASTLA, OASTLB and OASTLC level in WT and the various mutants analysed by Western blot employing specific antibody (A), total Se (B), selenate (C) and transcripts analyses of *OASTLA* (D), *OASTLB* (E) and *OASTLC* (F). Transcript levels of *OASTLA, OASTLB* and *OASTLC* were detected in the various genotypes by quantitative reverse transcription-PCR using *ACTIN2* as housekeeping transcript for normalization. The relative expression of each normalized *OASTL* in the various genotypes was analyzed using WT control as the reference. The values are means ± SE (n = 4). Values denoted with different letters are significantly different according to T-test analyses, P < 0.05 (JMP 8.0 software, http://www.jmp.com/). Different uppercase letters indicate significant differences between control and Se treatment of each genotype. Asterisks indicate significant differences between WT and *oas-tlA, oastlB, oastlC* KO plants subjected to the same treatment. The specific antibody kindly provided by Prof. Dr. S. Kopriva, University of Cologne, Germany.

**Supplemental Figure S3**. S/Se metabolism activities and DES1 transcript in WT and *Atdes1* KO mutants supplemented with or without selenate for 14 days. L-Cys/SeCys NATIVE-SDS degradation activities in the presence and absence of L-Cys/SeCys or BME (A), *DES1* transcripts expression with primers constructed upstream and downstream along the transcript length (B), OAS-TL (C) and DES kinetic activities (D), as well as, L-Cysteine (left side) and SeCys (right side) in-gel desulfhydrase activities in WT and *Atdes1* SALK_205358 leaves (E). DES1 transcript expression and in-gel L-Cysteine desulfhydrase activity in WT and *Atdes1* SALK_103855 (F). Transcript level of *Des1* was detected by quantitative reverse transcription-PCR using *ACTIN2* as the housekeeping transcript for normalization. The relative expression in the *Atdes1* with the 2 coupled primers was analyzed using the normalized WT control as reference. In-gel L-Cysteine desulfhydrase activities were employed after protein fractionation in SDS gel performed in Protean® II xi (www.bio-rad.com) in A and E and mini Protean3 (www.bio-rad.com) in F. The values are means ± SE (n = 4). Values denoted with different letters are significantly different according to T-test analyses, P < 0.05 (JMP 8.0 software, http://www.jmp.com/). Different uppercase letters indicate significant differences between control and Se treatment of each genotype. Different lower-case letters indicate differences between genotypes within the same primer. Asterisks indicate significant differences between WT and *des1* KO plants subjected to the same treatment

**Supplemental Figure S4**. Multiple sequence alignment of OASTL A, B and C proteins in *Arabidopsis thaliana*. The percentage of sequence identity between the proteins is presented below.

**Supplemental Figure S5**. Hydrogen peroxide (H_2_O_2_), malondialdehyde (MDA) and anthocyanin level in WT, *AtoastlA, AtoastlB* and *AtoastlC* KO mutants supplemented with or without selenate for 14 days. The effect of 40µM Selenate treatment on H_2_O_2_ (A), MDA (B) and anthocyanin (C) levels calculated on FW (left) and soluble protein basis (right). The values are means ± SE (n = 4). Values denoted with different letters are significantly different according to T-test analyses, P < 0.05 (JMP 8.0 software, http://www.jmp.com/). Different uppercase letters indicate significant differences between control and Se treatment of each genotype.

## Supplemental Tables

**Supplemental Table S1**. Identified and overlapped unique peptides from L-Cys and L-SeCys desulfhydrase activity. **A**. List of the number coverage percentage **B**. Identified and overlapped unique peptides from L-Cys desulfhydrase activity. **C**. Identified and overlapped unique peptides from L-SeCys desulfhydrase activity. D. Abundance of OASTL A, B and C as compared to DES1 protein at the lower, middle and upper activity bands. The semi quantitation was made by calculating the peak area of each peptide, whereas to overcome differential ionization, the area of the protein was calculated as the average of three most intense peptides from each protein. The calculated area of the DES protein is marked in red. The activity bands (as shown in Fig. 4A and B) were sliced and fractionated in 12.5 % SDS-PAGE. Further the bands were digested by trypsin and analyzed by LC-MS/MS as described in Materials and Methods. Blue color – overlapped areas. The presented data are representative of 3 independent experiments with similar results. *(see below).

**Supplemental Table S2**. List of gene primers used for quantitative real-time PCR.

## REFERENCES

Álvarez C, Calo L, Romero LC, García I, Gotor C (2010) An O-acetylserine (thiol) lyase homolog with L-cysteine desulfhydrase activity regulates cysteine homeostasis in Arabidopsis. Plant Physiology 152: 656–669

Barroso C, Vega J, Gotor C (1995) A new member of the cytosolic O-acetylserine(thiol)lyase gene family in Arabidopsis thaliana. FEBS Letters 363: 1–5

Bick J-A, Setterdahl AT, Knaff DB, Chen Y, Pitcher LH, Zilinskas BA, Leustek T (2001) Regulation of the plant-type 5 ‘-adenylyl sulfate reductase by oxidative stress. Biochemistry 40: 9040–9048

Bloem E, Riemenschneider A, Volker J, Papenbrock J, Schmidt A, Salac I, Haneklaus S, Schnug E (2004) Sulphur supply and infection with Pyrenopeziza brassicae influence L-cysteine desulphydrase activity in Brassica napus L. Journal of Experimental Botany 55: 2305–2312

Bradford MM (1976) A rapid and sensitive method for the quantitation of microgram quantities of protein utilizing the principle of protein-dye binding. Analytical biochemistry 72: 248–254

Brychkova G, Xia Z, Yang G, Yesbergenova Z, Zhang Z, Davydov O, Fluhr R, Sagi M (2007) Sulfite oxidase protects plants against sulfur dioxide toxicity. The Plant journal: for cell and molecular biology 50: 696–709

Brychkova G, Yarmolinsky D, Fluhr R, Sagi M (2012a) The determination of sulfite levels and its oxidation in plant leaves. Plant Sci 190: 123–130

Brychkova G, Yarmolinsky D, Ventura Y, Sagi M (2012b) A novel in-gel assay and an improved kinetic assay for determining in vitro sulfite reductase activity in plants. Plant and Cell Physiology 53: 1507–1516

Brychkova G, Grishkevich V, Fluhr R, Sagi M (2013) An essential role for tomato sulfite oxidase and enzymes of the sulfite network in maintaining leaf sulfite homeostasis. Plant physiology 161: 148–164

Brychkova G, Yarmolinsky D, Batushansky A, Grishkevich V, Khozin-Goldberg I, Fait A, Amir R, Fluhr R, Sagi M (2015) Sulfite oxidase activity is essential for normal sulfur, nitrogen and carbon metabolism in tomato leaves. Plants 4: 573–605

Domínguez-Solís JR, López-Martín MC, Ager FJ, Ynsa MD, Romero LC, Gotor C (2004) Increased cysteine availability is essential for cadmium tolerance and accumulation in Arabidopsis thaliana. Plant Biotechnology Journal 2: 469–476

Fisher B, Yarmolinsky D, Abdel-Ghany S, Pilon M, Pilon-Smits EA, Sagi M, Van Hoewyk D (2016) Superoxide generated from the glutathione-mediated reduction of selenite damages the iron-sulfur cluster of chloroplastic ferredoxin. Plant physiology and biochemistry 106: 228–235

Gaitonde M (1967) A spectrophotometric method for the direct determination of cysteine in the presence of other naturally occurring amino acids. Biochemical Journal 104: 627

Grant K, Carey NM, Mendoza M, Schulze J, Pilon M, Pilon-Smits EA, Van Hoewyk D (2011) Adenosine 5′-phosphosulfate reductase (APR2) mutation in Arabidopsis implicates glutathione deficiency in selenate toxicity. Biochemical Journal 438: 325–335

Gould KS, Mckelvie J, Markham KR (2002) Do anthocyanins function as antioxidants in leaves? Imaging of H2O2 in red and green leaves after mechanical injury. Plant Cell and Environment 25: 1261 – 1269

Gupta M, Gupta S (2017) An overview of selenium uptake, metabolism, and toxicity in plants. Frontiers in Plant Science 7: 2074

Heeg C, Kruse C, Jost R, Gutensohn M, Ruppert T, Wirtz M, Hell R (2008) Analysis of the Arabidopsis O-acetylserine (thiol) lyase gene family demonstrates compartment-specific differences in the regulation of cysteine synthesis. The Plant Cell 20: 168–185

Hesse H, Lipke J, Altmann T, Höfgen R (1999) Molecular cloning and expression analyses of mitochondrial and plastidic isoforms of cysteine synthase (O-acetylserine (thiol) lyase) from Arabidopsis thaliana. Amino Acids 16: 113–131

Hesse H, Hoefgen R (2003) Molecular aspects of methionine biosynthesis. Trends in plant science 8: 259–262

Holdorf MM, Owen HA, Lieber SR, Yuan L, Adams N, Dabney-Smith C, Makaroff CA (2012) Arabidopsis ETHE1 encodes a sulfur dioxygenase that is essential for embryo and endosperm development. Plant physiology 160: 226–236

Jin Z, Sun L, Yang G, Pei Y (2018) Hydrogen sulfide regulates energy production to delay leaf senescence induced by drought stress in Arabidopsis. Frontiers in Plant Science 9: 1722

Kalra Y (1997). “Handbook of reference methods for plant analysis,” CRC press.

Kant S, Kant P, Raveh E, Barak S (2006) Evidence that differential gene expression between the halophyte, Thellungiella halophila, and Arabidopsis thaliana is responsible for higher levels of the compatible osmolyte proline and tight control of Na+ uptake in T. halophila. Plant, Cell Environ 29: 1220–1234

Khan MS, Haas FH, Samami AA, Gholami AM, Bauer A, Fellenberg K, Reichelt M, Hänsch R, Mendel RR, Meyer AJ (2010) Sulfite reductase defines a newly discovered bottleneck for assimilatory sulfate reduction and is essential for growth and development in Arabidopsis thaliana. The Plant Cell 22: 1216–1231

Koprivova A, North KA, Kopriva S (2008) Complex signaling network in regulation of adenosine 5′-phosphosulfate reductase by salt stress in Arabidopsis roots. Plant Physiology 146: 1408–1420

Krüßel L, Junemann J, Wirtz M, Birke H, Thornton JD, Browning LW, Poschet G, Hell R, Balk J, Braun H-P (2014) The mitochondrial sulfur dioxygenase ETHYLMALONIC ENCEPHALOPATHY PROTEIN1 is required for amino acid catabolism during carbohydrate starvation and embryo development in Arabidopsis. Plant physiology 165: 92–104

Kurmanbayeva A, Bekturova A, Srivastava S, Soltabayeva A, Asatryan A, Ventura Y, Khan MS, Salazar O, Fedoroff N, Sagi M (2017) Higher novel L-Cys degradation activity results in lower organic-S and biomass in Sarcocornia than the related saltwort, Salicornia. Plant physiology 175: 272–289

Kuske CR, Hill KK, Guzman E, Jackson PJ (1996) Subcellular location of O-acetylserine sulfhydrylase isoenzymes in cell cultures and plant tissues of Datura innoxia Mill. Plant physiology 112: 659–667

Laby RJ, Kincaid MS, Kim D, Gibson SI (2000) The Arabidopsis sugar-insensitive mutants sis4 and sis5 are defective in abscisic acid synthesis and response. Plant J 23: 587–596

Lang C, Popko J, Wirtz M, Hell R, Herschbach C, Kreuzwieser J, Rennenberg H, Mendel RR, Haensch R (2007) Sulphite oxidase as key enzyme for protecting plants against sulphur dioxide. Plant, Cell & Environment 30: 447–455

Laemmli UK (1970) Cleavage of structural proteins during the assembly of the head of bacteriophage T4. nature 227: 680

Lee BR, Koprivova A, Kopriva S (2011) The key enzyme of sulfate assimilation, adenosine 5′-phosphosulfate reductase, is regulated by HY5 in Arabidopsis. The Plant Journal 67: 1042–1054

Lima LW, Pilon-Smits EA, Schiavon M (2018) Mechanisms of selenium hyperaccumulation in plants: A survey of molecular, biochemical and ecological cues. Biochimica et Biophysica Acta (BBA)-General Subjects 1862: 2343–2353

Manchenko GP (2002). “Handbook of detection of enzymes on electrophoretic gels,” CRC press.

Mroczek-Zdyrska M, Wójcik M (2012) The influence of selenium on root growth and oxidative stress induced by lead in Vicia faba L. minor plants. Biological trace element research 147: 320–328

Murashige T, Skoog F (1962) A revised medium for rapid growth and bio assays with tobacco tissue cultures. Physiologia plantarum 15: 473–497

Papp LV, Lu J, Holmgren A, Khanna KK (2007) From selenium to selenoproteins: synthesis, identity, and their role in human health. Antioxidants & redox signaling 9: 775–806

Pilon-Smits EA, Hwang S, Lytle CM, Zhu Y, Tai JC, Bravo RC, Chen Y, Leustek T, Terry N (1999) Overexpression of ATP sulfurylase in Indian mustard leads to increased selenate uptake, reduction, and tolerance. Plant Physiology 119: 123–132

Pilon-Smits EA, Quinn CF (2010). Selenium metabolism in plants. In “Cell biology of metals and nutrients”, pp. 225–241. Springer.

Pilon-Smits EA, Womg M (2012). Plant selenium metabolism–genetic manipulation, phytotechnological applications, and ecological implications. In “Environmental contamination: Health risks and ecological restoration”, pp. 293–311. CRC Press Boca Raton, FL.

Ravanel S, Gakière B, Job D, Douce R (1998) The specific features of methionine biosynthesis and metabolism in plants. Proceedings of the National Academy of Sciences 95: 7805–7812

Riemenschneider A, Riedel K, Hoefgen R, Papenbrock J, Hesse H (2005) Impact of reduced O-acetylserine (thiol) lyase isoform contents on potato plant metabolism. Plant physiology 137: 892–900

Ritchie RJ (2006) Consistent sets of spectrophotometric chlorophyll equations for acetone, methanol and ethanol solvents. Photosynth Res 89: 27–41

Sabbagh M, Van Hoewyk D (2012) Malformed selenoproteins are removed by the ubiquitin– proteasome pathway in Stanleya pinnata. Plant and Cell Physiology 53: 555–564

Sagi M, Fluhr R (2001) Superoxide production by plant homologues of the gp91phox NADPH oxidase. Modulation of activity by calcium and by tobacco mosaic virus infection. Plant Physiology 126: 1281–1290

Schiavon M, Moro I, Pilon-Smits EA, Matozzo V, Malagoli M, Dalla Vecchia F (2012) Accumulation of selenium in Ulva sp. and effects on morphology, ultrastructure and antioxidant enzymes and metabolites. Aquatic toxicology 122: 222–231

Schiavon M, Pilon M, Malagoli M, Pilon-Smits EA (2015) Exploring the importance of sulfate transporters and ATP sulphurylases for selenium hyperaccumulation—a comparison of Stanleya pinnata and Brassica juncea (Brassicaceae). Frontiers in plant science 6: 2

Seko Y, Saito Y, Kitahara J, Imura N (1989). Active oxygen generation by the reaction of selenite with reduced glutathione in vitro. In “Selenium in biology and medicine”, pp. 70–73. Springer.

Siegel LM (1965) A direct microdetermination for sulfide. Analytical biochemistry 11: 126–132

Sors TG, Ellis DR, Na GN, Lahner B, Lee S, Leustek T, Pickering IJ, Salt DE (2005) Analysis of sulfur and selenium assimilation in Astragalus plants with varying capacities to accumulate selenium. The Plant Journal 42: 785–797

Srivastava S, Brychkova G, Yarmolinsky D, Soltabayeva A, Samani T, Sagi M (2017) Aldehyde Oxidase 4 plays a critical role in delaying silique senescence by catalyzing aldehyde detoxification. Plant physiology 173: 1977–1997

Terry N, Zayed A, De Souza M, Tarun A (2000) Selenium in higher plants. Annual review of plant biology 51: 401–432

Tian M, Hui M, Thannhauser TW, Pan S, Li L (2017) Selenium-induced toxicity is counteracted by sulfur in broccoli (Brassica oleracea L. var. italica). Frontiers in plant science 8: 1425

Van Hoewyk D (2018) Defects in endoplasmic reticulum-associated degradation (ERAD) increase selenate sensitivity in Arabidopsis. Plant signaling & behavior 13: e1171451

Van Hoewyk D, Garifullina GF, Ackley AR, Abdel-Ghany SE, Marcus MA, Fakra S, Ishiyama K, Inoue E, Pilon M, Takahashi H (2005) Overexpression of AtCpNifS enhances selenium tolerance and accumulation in Arabidopsis. Plant Physiology 139: 1518–1528

White PJ (2015) Selenium accumulation by plants. Annals of botany 117: 217–235

White PJ (2018) Selenium metabolism in plants. Biochimica et Biophysica Acta (BBA)-General Subjects 1862: 2333–2342

White PJ, Bowen HC, Parmaguru P, Fritz M, Spracklen W, Spiby R, Meacham M, Mead A, Harriman M, Trueman L (2004) Interactions between selenium and sulphur nutrition in Arabidopsis thaliana. Journal of Experimental Botany 55: 1927–1937

Wirtz M, Hell R (2006) Functional analysis of the cysteine synthase protein complex from plants: structural, biochemical and regulatory properties. Journal of plant physiology 163: 273–286

Xu Z, Mahmood K, Rothstein SJ (2017) ROS induces anthocyanin production via late biosynthetic genes and anthocyanin deficiency confers the hypersensitivity to ROS-generating stresses in Arabidopsis. Plant and Cell Physiology 58: 1364–1377

Yarmolinsky D, Brychkova G, Kurmanbayeva A, Bekturova A, Ventura Y, Khozin-Goldberg I, Eppel A, Fluhr R, Sagi M (2014) Impairment in Sulfite Reductase Leads to Early Leaf Senescence in Tomato Plants. Plant Physiology 165: 1505–1520

Yarmolinsky D, Brychkova G, Fluhr R, Sagi M (2013) Sulfite reductase protects plants against sulfite toxicity. Plant physiology 161: 725–743

Yesbergenova Z, Yang G, Oron E, Soffer D, Fluhr R, Sagi M (2005) The plant Mo-hydroxylases aldehyde oxidase and xanthine dehydrogenase have distinct reactive oxygen species signatures and are induced by drought and abscisic acid. The Plant Journal 42: 862–876

