## Supplementary material for "Impairment in O-acetylserine-(thiol) lyase A and B, but not C, confers higher selenate sensitivity and uncovers role for A, B and C as L-Cys and L-SeCys desulfhydrases in *Arabidopsis*": Supp. Figures 1-6 and Tables 1-2

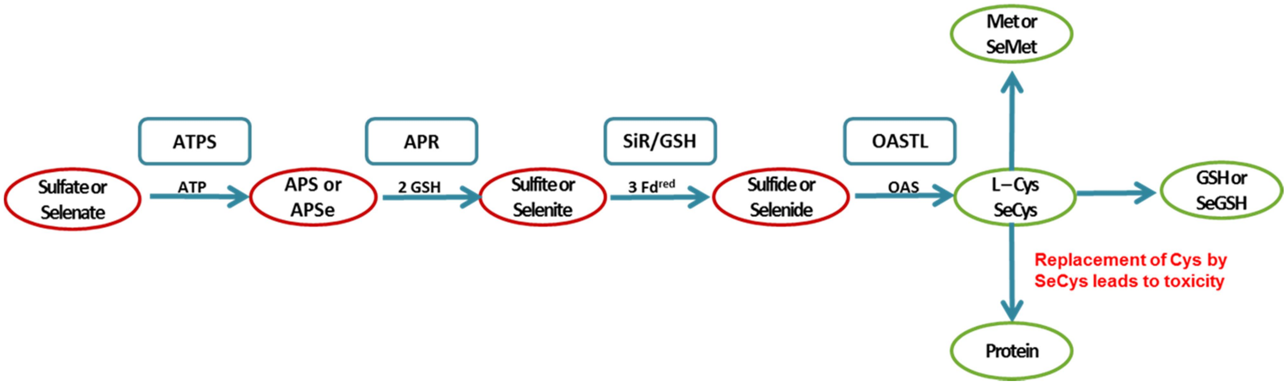


**Figure S1.** Schematic model of Sulfur (S) and Selenium (Se) metabolism in Arabidopsis plants [after White (2015)]. ATP sufurylase (ATPS) catalyzes the adenylation of sulfate/selenate to adenosine 5′-phosphosulfate (APS)/adenosine phosphoselenate (APSe) using ATP as an electron donor. Then, APS/APSe is reduced by the plastidic enzyme APS reductase (APR) to sulfite/selenite in the presence of two molecules of reduced glutathione, which acts as an electron donor. The generated sulfite/selenite can be reduced to sulfide/selenide by the Sulfite Reductase (SiR) employing 3 molecules of reduced ferredoxin, alternatively, selenite also can reduced non-enzymatically by glutathione (GSH) to selenide. The sulfide/selenide together with O-acetyl-L-Serine (OAS) are the substrates for Cysteine (Cys)/Selenocysteine (SeCys) biosynthesis catalyzed by O-acetylserine-(thiol) lyase (OAS-TL).

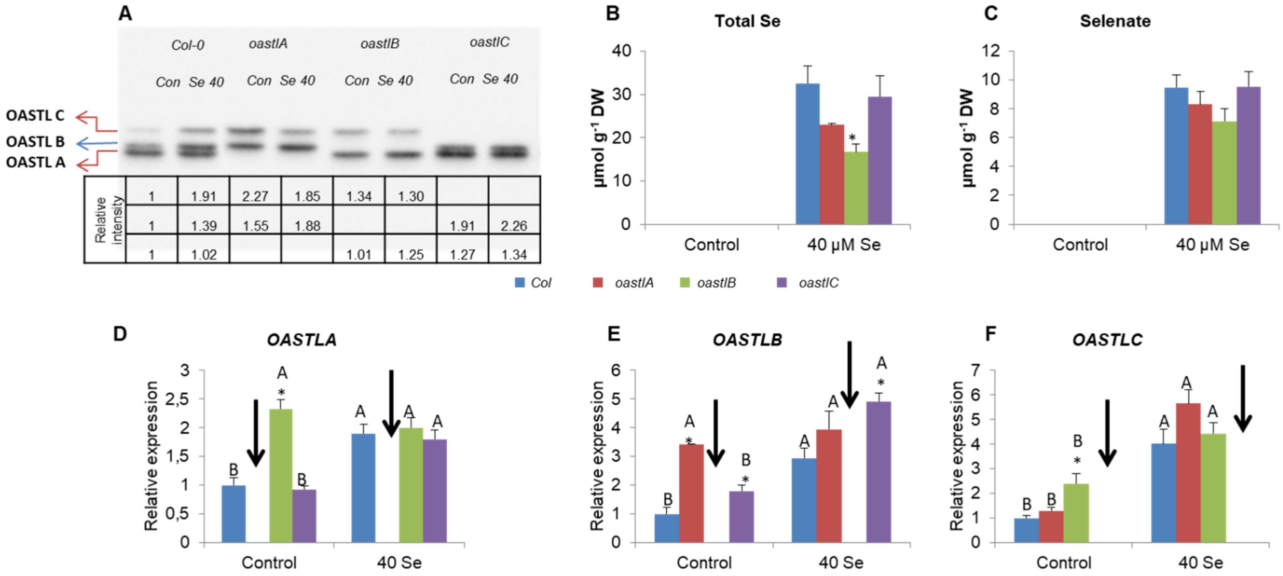


**Supplemental Figure S2.** OASTLs protein and transcripts expression, as well as Se containing compounds in *WT, AtoastlA, AtoastlB* and *AtoastlC* KO mutants supplemented with or without selenate for 14 days. Effect of 40µM Selenate treatment on OASTLA, OASTLB and OASTLC level in WT and the various mutants analysed by Western blot employing specific antibody (A), total Se (B), selenate (C) and transcripts analyses of *OASTLA* (D), *OASTLB* (E) and *OASTLC* (F). Transcript levels of *OASTLA, OASTLB* and *OASTLC* were detected in the various genotypes by quantitative reverse transcription-PCR using *ACTIN2* as housekeeping transcript for normalization. The relative expression of each normalized *OASTL* in the various genotypes was analyzed using WT control as the reference. The values are means ± SE (n = 4). Values denoted with different letters are significantly different according to T-test analyses, P < 0.05 (JMP 8.0 software, http://www.jmp.com/). Different uppercase letters indicate significant differences between control and Se treatment of each genotype. Asterisks indicate significant differences between WT and *oas-tlA, oastlB, oastlC* KO plants subjected to the same treatment. The specific antibody was kindly provided by Prof. Dr. S. Kopriva, University of Cologne, Germany.

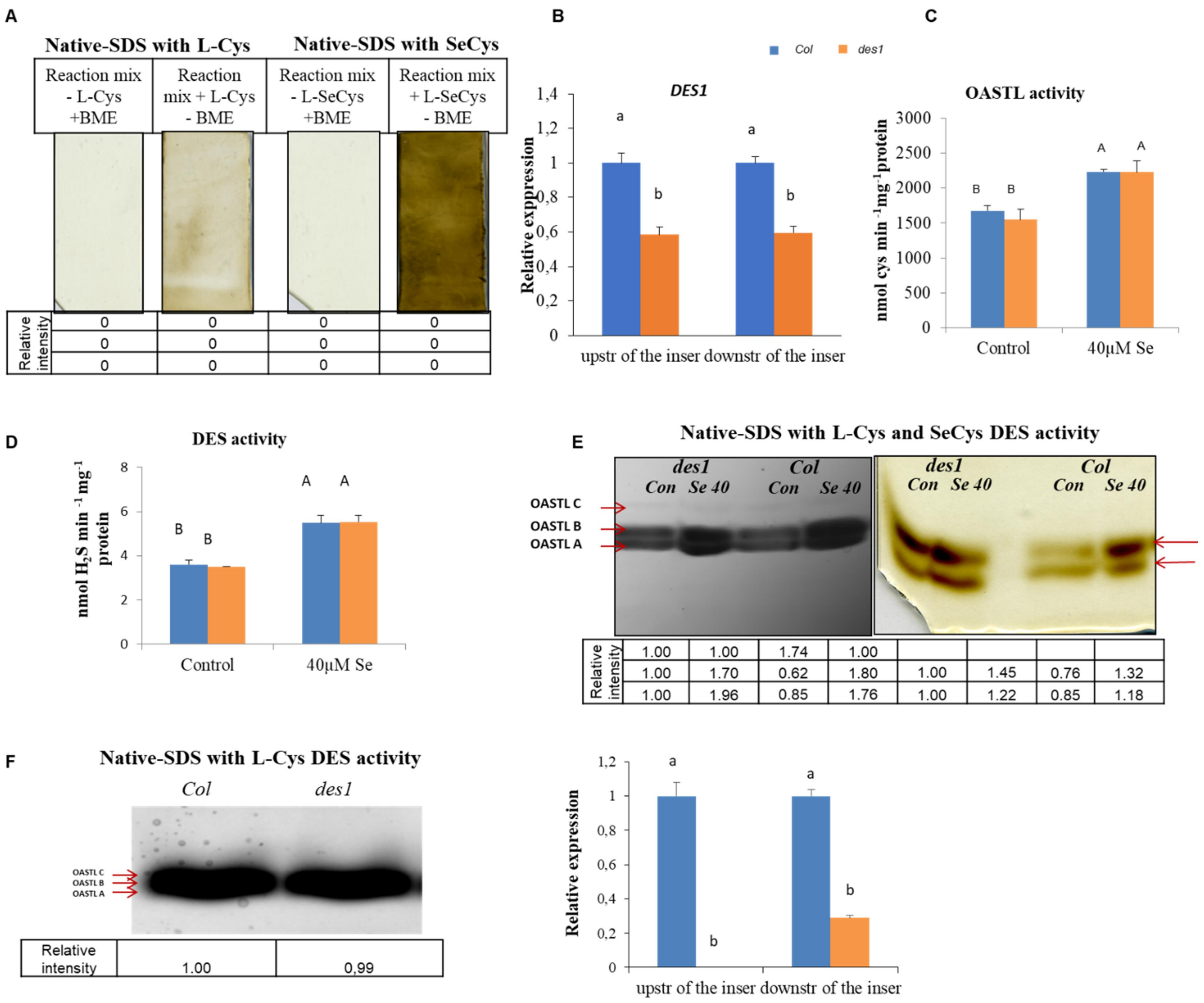


**Supplemental Figure S3.** S/Se metabolism activities and DES1 transcript in WT and *Atdes1* KO mutants supplemented with or without selenate for 14 days. L-Cys/SeCys NATIVE-SDS degradation activities in the presence and absence of L-Cys/SeCys or BME (A), *DES1* transcripts expression with primers constructed upstream and downstream along the transcript length (B), OAS-TL (C) and DES kinetic activities (D), as well as, L-Cysteine (left side) and SeCys (right side) in-gel desulfhydrase activities in WT and *Atdes1* SALK_205358 leaves (E). DES1 transcript expression and in-gel L-Cysteine desulfhydrase activity in WT and *Atdes1* SALK_103855 (F). Transcript level of *Des1* was detected by quantitative reverse transcription-PCR using *ACTIN2* as the housekeeping transcript for normalization. The relative expression in the *Atdes1* with the 2 coupled primers was analyzed using the normalized WT control as reference. In-gel L-Cysteine desulfhydrase activities were employed after protein fractionation in SDS gel performed in Protean® II xi (www.bio-rad.com) in A and E and mini Protean3 (www.bio-rad.com) in F. The values are means ± SE (n = 4). Values denoted with different letters are significantly different according to T-test analyses, $P < 0.05$ (JMP 8.0 software, http://www.jmp.com/). Different uppercase letters indicate significant differences between control and Se treatment of each genotype. Different lower-case letters indicate differences between genotypes within the same primer. Asterisks indicate significant differences between WT and *des1* KO plants subjected to the same treatment

```
OASTLA      ------------------------------------------------------------    0
OASTLB      --------------------------------MAATSSSAFLLNPLTSR-----HRPFK    22
OASTLC      MVAMIMASRFNREAKLASQILSTLLGNRSCYTSMAATSSSALLLNPLTSSSSSSTLRRFR    60


OASTLA      --------------------------------------------------MASRIAKDVT    10
OASTLB      YSPELSSLSLSSRKAAAFDVSS--AAFTLKRQSRSDVVCKAVSIKPEAGVEGLNIADNAA    80
OASTLC      CSPEISSLSFSSASDFSLAMKRQSRSFADGSERDPSVVCEA--VKRETGPDGLNIADNVS    118
                                                               . .**.:.:


OASTLA      ELIGNTPLVYLNNVAEGCVGRVAAKLEMMEPCSSVKDRIGFSMISDAEKKGLIKPGESVL    70
OASTLB      QLIGKTPMVYLNNVVKGCVASVAAKLEIMEPCCSVKDRIGYSMITDAEEKGLITPGKSVL    140
OASTLC      QLIGKTPMVYLNSIAKGCVANIAAKLEIMEPCCSVKDRIGYSMVTDAEQKGFISPGKSVL    178
            :***:**:****.:.:***. :*****:****.*******:**::***:**:*.**:***


OASTLA      IEPTSGNTGVGLAFTAAAKGYKLIITMPASMSTERRIILLAFGVELVLTDPAKGMKGAIA    130
OASTLB      VESTSGNTGIGLAFIAASKGYKLILTMPASMSLERRVLLRAFGAELVLTEPAKGMTGAIQ    200
OASTLC      VEPTSGNTGIGLAFIAASRGYRLILTMPASMSMERRVLLKAFGAELVLTDPAKGMTGAVQ    238
            :* ******:**** **::**:**:******* ***::* ***.*****:*****.**:


OASTLA      KAEEILAKTPNGYMLQQFENPANPKIHYETTGPEIWKGTGGKIDGFVSGIGTGGTITGAG    190
OASTLB      KAEEILKKTPNSYMLQQFDNPANPKIHYETTGPEIWEDTRGKIDILVAGIGTGGTITGVG    260
OASTLC      KAEEILKNTPDAYMLQQFDNPANPKIHYETTGPEIWDDTKGKVDIFVAGIGTGGTITGVG    298
            ****** :**:.******:*****************..* **:* :*:**********.*

      1: OASTLA      100.00   71.12   70.19
      2: OASTLB       71.12  100.00   79.74
      3: OASTLC       70.19   79.74  100.00**
```

**Supplemental Figure S4.** Multiple sequence alignment of OASTL A, B and C proteins of in *Arabidopsis thaliana*. **The percentage of sequence identity between the proteins is presented below.

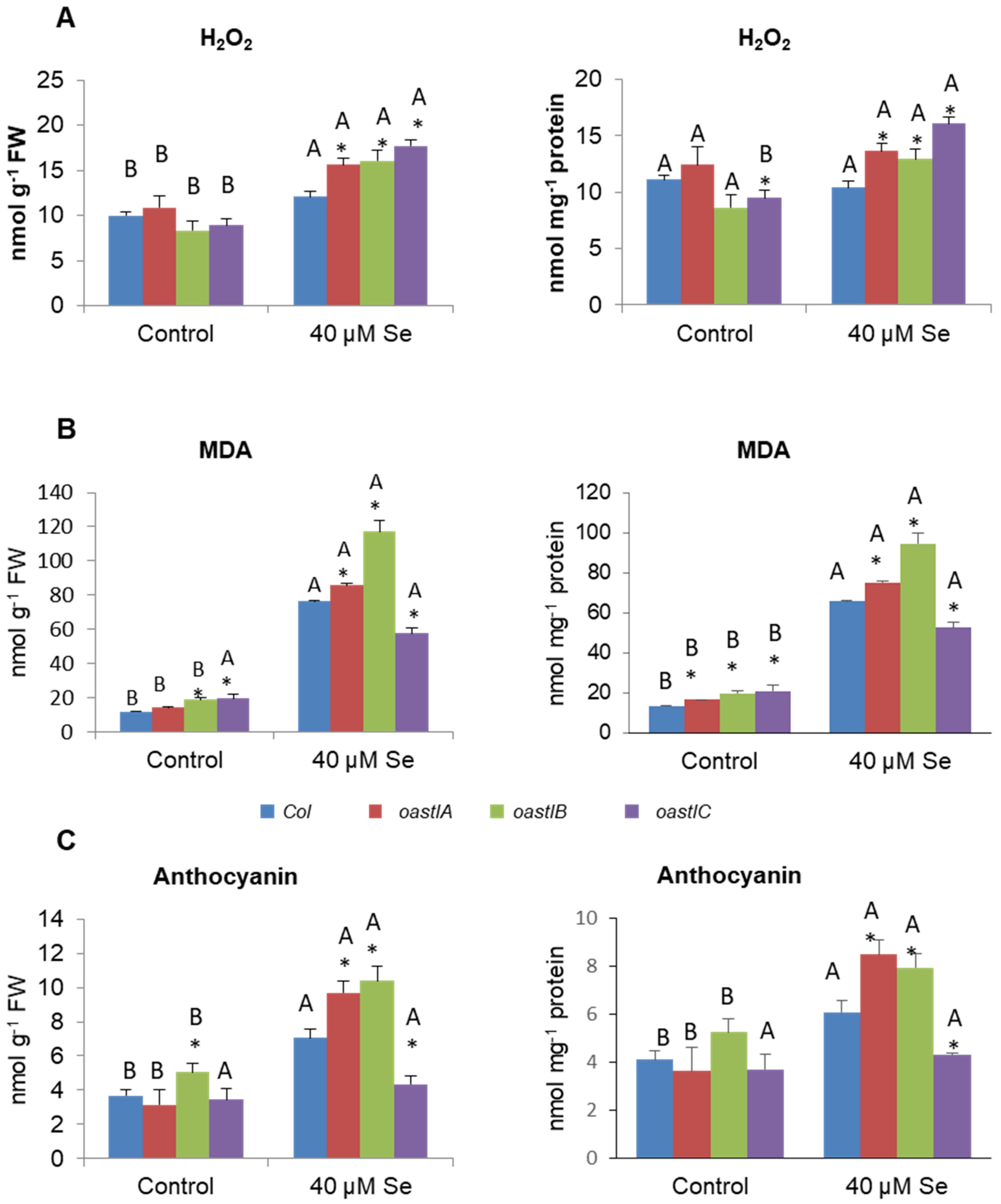


**Supplemental Figure S5.** Hydrogen peroxide ($H_2O_2$), malondialdehyde (MDA) and anthocyanin level in WT, *AtoastlA, AtoastlB* and *AtoastlC* KO mutants supplemented with or without selenate for 14 days. The effect of 40µM Selenate treatment on $H_2O_2$ (A), MDA (B) and anthocyanin (C) levels calculated on FW (left) and soluble protein basis (right). The values are means ± SE (n = 4). Values denoted with different letters are significantly different according to T-test analyses, $P < 0.05$ (JMP 8.0 software, http://www.jmp.com/). Different uppercase letters indicate significant differences between control and Se treatment of each genotype.

**Supplemental Table S1.** Identified and overlapped unique peptides from L-Cys and L-SeCys desulfhydrase activity. **A**. List of the number coverage percentage **B**. Identified and overlapped unique peptides from L-Cys desulfhydrase activity. **C**. Identified and overlapped unique peptides from L-SeCys desulfhydrase activity. D. Abundancy of OASTL A, B and C as compared to DES1 protein in the lower, middle and upper activity bands. The semi quantitation was made by calculating the peak area of each peptide, whereas to overcome differential ionization, the area of the protein was calculated as the average of three most intense peptides from each protein. The calculated area of the DES protein is marked in red. The activity bands (as shown in Fig. 4A and B) were sliced and fractionated in 12.5 % SDS-PAGE. Further the bands were digested by trypsin and analyzed by LC-MS/MS as described in Materials and Methods. Blue color – overlapped areas. The presented data are representative of 3 independent experiments with similar results. *(see below).

**A**

| With L-Cys | OASTL A P47998 | OASTL B P47999 | OASTL C Q43725 | DES1 F4K5T2 |
|---|---|---|---|---|
| **Col upper band** | 6peptides<br>71aa<br>22% | 11peptides<br>136aa<br>35% | 13peptides<br>193aa<br>45% | |
| **Col middle band** | 18peptides<br>184aa<br>57% | 23peptides<br>227aa<br>58% | 6peptides<br>99aa<br>23% | 2peptides<br>22aa<br>6.8% |
| **Col Lower band** | 23peptides<br>213aa<br>66% | 14peptides<br>150aa<br>38% | | 3peptides<br>25aa<br>7.7% |

| With SeCys | OASTLA | OASTLB | OASTLC | DES1 |
|---|---|---|---|---|
| **Col middle band** | 15peptides<br>177aa<br>55% | 18peptides<br>237aa<br>60.5% | | |
| **Col Lower band** | 20peptides<br>208aa<br>65% | 9peptides<br>116aa<br>29.6% | | |

## B

| | | | | |
|---|---|---|---|---|
| Col upper band with L-Cys | **P47998.2**<br>**OAS-TL A - 6 peptides**<br>**71 aa are covered from 322 aa - 22%** | **P47999.2**<br>**OAS-TL B – 11peptides**<br>**136 aa are covered from 392 aa 38 %** | **Q43725**<br>**OASTLC – 13 peptides**<br>**193 aa are covered from 430 aa 45 %** | **F4K5T2**<br>**Des1 – 0** |
| | MASRIAKDVTELIGNTPLVYLNNVAEGCVGRVAAKLE<br>MMEPCSSVKDRIGFSMISDAEKKGLIKPGESVLIEPTS<br>GNTGVGLAFTAAAKGYKLIITMPASMSTERRIILLAFG<br>VELVLTDPAKGMKGAIAKAEEILAKTPNGYMLQQFE<br>NPANPKIHYETTGPEIWKGTGGKIDGFVSGIGTGGTI<br>TGAGKYLKEQNANVKLYGVEPVESAILSGGKPGPHKI<br>QGIGAGFIPSVLNVDLIDEVVQVSSDESIDMARQLAL<br>KEGLLVGISSGAAAAAAIKLAQRPENAGKLFVAIFPSF<br>GERYLSTVLFDATRKEAEAMTFEA 322aa covered<br>71aa 22% | MAATSSSAFLLNPLTSRHRPFKYSPELSSLSLSSRKAAAFDV<br>SSAAFTLKRQSRSDVVCKAVSIKPEAGVEGLNIADNAAQLI<br>GKTPMVYLNNVVKGCVASVAAKLEIMEPCCSVKDRIGYS<br>MITDAEEKGLITPGKSVLVESTSGNTGIGLAFIAASKGYKLIL<br>TMPASMSLERRVLLRAFGAELVLTEPAKGMTGAIQKAEEI<br>LKKTPNSYMLQQFDNPANPKIHYETTGPEIWEDTRGKIDIL<br>VAGIGTGGTITGVGRFIKERKPELKVIGVEPTESAILSGGKP<br>GPHKIQGIGAGFVPKNLDLAIVDEYIAISSEEAIETSKQLALQ<br>EGLLVGISSGAAAAAAIQVAKRPENAGKLIAVVFPSFGERY<br>LSTQLFQSIREECEQMQPEL | MVAMIMASRFNREAKLASQILSTLLGNRSCYTSMAATSSSALLL<br>NPLTSSSSSSTLRRFRCSPEISSLSFSSASDFSLAMKRQSRSFADG<br>SERDPSVVCEAVKRETGPDGLNIADNVSQLIGKTPMVYLNSIAK<br>GCVANIAAKLEIMEPCCSVKDRIGYSMVTDAEQKGFISPGKSVL<br>VEPTSGNTGIGLAFIAASRGYRLILTMPASMSMERRVLLKAFGA<br>ELVLTDPAKGMTGAVQKAEEILKNTPDAYMLQQFDNPANPKI<br>HYETTGPEIWDDTKGKVDIFVAGIGTGGTITGVGRFIKEKNPKT<br>QVIGVEPTESDILSGGKPGPHKIQGIGAGFIPKNLDQKIMDEVIA<br>ISSEEAIETAKQLALKEGLMVGISSGAAAAAAIKVAKRPENAGKL<br>IAVVFPSFGERYLSTPLFQSIREEVEKMQPEV | |
| Col Mid band with L-Cys | **P47998.2**<br>**OAS-TL A - 18 peptides**<br>**184 aa are covered from 322 aa - 57%** | **P47999.2**<br>**OAS-TL B – 23 peptides**<br>**227 aa are covered from 392 aa 58 %** | **Q43725**<br>**OASTLC – 6 peptides**<br>**99 aa are covered from 430 aa 23 %** | **F4K5T2**<br>**Des1 –2peptides**<br>**22 aa are covered from 323 aa – 6.8%** |
| | MASRIAKDVTELIGNTPLVYLNNVAEGCVGRVAAKLE<br>MMEPCSSVKDRIGFSMISDAEKKGLIKPGESVLIEPTS<br>GNTGVGLAFTAAAKGYKLIITMPASMSTERRIILLAFG<br>VELVLTDPAKGMKGAIAKAEEILAKTPNGYMLQQFE<br>NPANPKIHYETTGPEIWKGTGGKIDGFVSGIGTGGTI<br>TGAGKYLKEQNANVKLYGVEPVESAILSGGKPGPHKI<br>QGIGAGFIPSVLNVDLIDEVVQVSSDESIDMARQLAL<br>KEGLLVGISSGAAAAAAIKLAQRPENAGKLFVAIFPSF<br>GERYLSTVLFDATRKEAEAMTFEA | MAATSSSAFLLNPLTSRHRPFKYSPELSSLSLSSRKAAAFDV<br>SSAAFTLKRQSRSDVVCKAVSIKPEAGVEGLNIADNAAQLI<br>GKTPMVYLNNVVKGCVASVAAKLEIMEPCCSVKDRIGYS<br>MITDAEEKGLITPGKSVLVESTSGNTGIGLAFIAASKGYKLIL<br>TMPASMSLERRVLLRAFGAELVLTEPAKGMTGAIQKAEEI<br>LKKTPNSYMLQQFDNPANPKIHYETTGPEIWEDTRGKIDIL<br>VAGIGTGGTITGVGRFIKERKPELKVIGVEPTESAILSGGKP<br>GPHKIQGIGAGFVPKNLDLAIVDEYIAISSEEAIETSKQLALQ<br>EGLLVGISSGAAAAAAIQVAKRPENAGKLIAVVFPSFGERY<br>LSTQLFQSIREECEQMQPEL | MVAMIMASRFNREAKLASQILSTLLGNRSCYTSMAATSSSALLL<br>NPLTSSSSSSTLRRFRCSPEISSLSFSSASDFSLAMKRQSRSFADG<br>SERDPSVVCEAVKRETGPDGLNIADNVSQLIGKTPMVYLNSIAK<br>GCVANIAAKLEIMEPCCSVKDRIGYSMVTDAEQKGFISPGKSVL<br>VEPTSGNTGIGLAFIAASRGYRLILTMPASMSMERRVLLKAFGA<br>ELVLTDPAKGMTGAVQKAEEILKNTPDAYMLQQFDNPANPKI<br>HYETTGPEIWDDTKGKVDIFVAGIGTGGTITGVGRFIKEKNPKT<br>QVIGVEPTESDILSGGKPGPHKIQGIGAGFIPKNLDQKIMDEVIA<br>ISSEEAIETAKQLALKEGLMVGISSGAAAAAAIKVAKRPENAGKL<br>IAVVFPSFGERYLSTPLFQSIREEVEKMQPEV | MEDRVLIKNDVTELIGNTPMVYLNKIVDGCV<br>ARIAAKLEMMEPCSSIKDRIAYSMIKDAEDK<br>GLITPGKSTLIEATGGNTGIGLASIGASRGYKV<br>ILLMPSTMSLERRIILRALGAEVHLTDISIGIKG<br>QLEKAKEILSKTPGGYIPHQFINPENPEIHYRT<br>TGPEIWRDSAGKVDILVAGVGTGGTVTGTG<br>KFLKEKNKDIKVCVVEPSESAVLSGGKPGPHL<br>IQGIGSGEIPANLDLSIVDEIIQVTGEEAIETTK<br>LLAIKEGLLVGISSGASAAAALKVAKRPENVG<br>KLIVVIFPSGGERYLSTELFESVRYEAENLPVE |
| Col Lower band with L-Cys | **P47998.2**<br>**OAS-TL A - 23 peptides**<br>**213 aa are covered from 322 aa - 66%** | **P47999.2**<br>**OAS-TL B – 14 peptides**<br>**150 aa are covered from 392 aa 38 %** | **Q43725**<br>**OASTLC – 0** | **F4K5T2**<br>**Des1 –3peptides**<br>**25 aa are covered from 323 aa -7.7%** |
| | MASRIAKDVTELIGNTPLVYLNNVAEGCVGRVAAKLE<br>MMEPCSSVKDRIGFSMISDAEKKGLIKPGESVLIEPTS<br>GNTGVGLAFTAAAKGYKLIITMPASMSTERRIILLAFG<br>VELVLTDPAKGMKGAIAKAEEILAKTPNGYMLQQFE<br>NPANPKIHYETTGPEIWKGTGGKIDGFVSGIGTGGTI<br>TGAGKYLKEQNANVKLYGVEPVESAILSGGKPGPHKI<br>QGIGAGFIPSVLNVDLIDEVVQVSSDESIDMARQLAL<br>KEGLLVGISSGAAAAAAIKLAQRPENAGKLFVAIFPSF<br>GERYLSTVLFDATRKEAEAMTFEA | MAATSSSAFLLNPLTSRHRPFKYSPELSSLSLSSRKAAAFDV<br>SSAAFTLKRQSRSDVVCKAVSIKPEAGVEGLNIADNAAQLI<br>GKTPMVYLNNVVKGCVASVAAKLEIMEPCCSVKDRIGYS<br>MITDAEEKGLITPGKSVLVESTSGNTGIGLAFIAASKGYKLIL<br>TMPASMSLERRVLLRAFGAELVLTEPAKGMTGAIQKAEEI<br>LKKTPNSYMLQQFDNPANPKIHYETTGPEIWEDTRGKIDIL<br>VAGIGTGGTITGVGRFIKERKPELKVIGVEPTESAILSGGKP<br>GPHKIQGIGAGFVPKNLDLAIVDEYIAISSEEAIETSKQLALQ<br>EGLLVGISSGAAAAAAIQVAKRPENAGKLIAVVFPSFGERY<br>LSTQLFQSIREECEQMQPEL | | MEDRVLIKNDVTELIGNTPMVYLNKIVDGCV<br>ARIAAKLEMMEPCSSIKDRIAYSMIKDAEDK<br>GLITPGKSTLIEATGGNTGIGLASIGASRGYKV<br>ILLMPSTMSLERRIILRALGAEVHLTDISIGIKG<br>QLEKAKEILSKTPGGYIPHQFINPENPEIHYRT<br>TGPEIWRDSAGKVDILVAGVGTGGTVTGTG<br>KFLKEKNKDIKVCVVEPSESAVLSGGKPGPHL<br>IQGIGSGEIPANLDLSIVDEIIQVTGEEAIETTK<br>LLAIKEGLLVGISSGASAAAALKVAKRPENVG<br>KLIVVIFPSGGERYLSTELFESVRYEAENLPVE |

## C

| | | | | |
|---|---|---|---|---|
| Col Mid band with SeCys | **P47998.2**<br>**OAS-TL A - 15 peptides**<br>**177 aa are covered from 322 aa - 55%** | **P47999.2**<br>**OAS-TL B – 18 peptides**<br>**237aa are covered from 392 aa 60.5 %** | **Q43725 OASTLC – 0** | **F4K5T2 Des1 – 0** |
| | MASRIAKDVTELIGNTPLVYLNNVAEGCVGRVAAKLEMMEPCSSVKDRIGFSMISDAE<br>KKGLIKPGESVLIEPTSGNTGVGLAFTAAAKGYKLIITMPASMSTERRIILLAFGVELVLTD<br>PAKGMKGAIAKAEEILAKTPNGYMLQQFENPANPKIHYETTGPEIWKGTGGKIDGFVS<br>GIGTGGTITGAGKYLKEQNANVKLYGVEPVESAILSGGKPGPHKIQGIGAGFIPSVLNVD<br>LIDEVVQVSSDESIDMARQLALKEGLLVGISSGAAAAAAIKLAQRPENAGKLFVAIFPSF<br>GERYLSTVLFDATRKEAEAMTFEA | MAATSSSAFLLNPLTSRHRPFKYSPELSSLSLSSRKAAAFDVSSAAFTLKRQSRSDVVCKAVSIK<br>PEAGVEGLNIADNAAQLIGKTPMVYLNNVVKGCVASVAAKLEIMEPCCSVKDRIGYSMITDA<br>EEKGLITPGKSVLVESTSGNTGIGLAFIAASKGYKLILTMPASMSLERRVLLRAFGAELVLTEPA<br>KGMTGAIQKAEEILKKTPNSYMLQQFDNPANPKIHYETTGPEIWEDTRGKIDILVAGIGTGGT<br>ITGVGRFIKERKPELKVIGVEPTESAILSGGKPGPHKIQGIGAGFVPKNLDLAIVDEYIAISSEEAI<br>ETSKQLALQEGLLVGISSGAAAAAAIQVAKRPENAGKLIAVVFPSFGERYLSTQLFQSIREECE<br>QMQPEL | | |
| Col Lower band with SeCys | **P47998.2**<br>**OAS-TL A - 20 peptides**<br>**208 aa are covered from 322 aa - 65%** | **P47999.2**<br>**OAS-TL B – 9 peptides**<br>**116 aa are covered from 392 aa 29.6 %** | **Q43725 OASTLC – 0** | **F4K5T2 Des1 – 0** |
| | MASRIAKDVTELIGNTPLVYLNNVAEGCVGRVAAKLEMMEPCSSVKDRIGFSMISDAE<br>KKGLIKPGESVLIEPTSGNTGVGLAFTAAAKGYKLIITMPASMSTERRIILLAFGVELVLTD<br>PAKGMKGAIAKAEEILAKTPNGYMLQQFENPANPKIHYETTGPEIWKGTGGKIDGFVS<br>GIGTGGTITGAGKYLKEQNANVKLYGVEPVESAILSGGKPGPHKIQGIGAGFIPSVLNVD<br>LIDEVVQVSSDESIDMARQLALKEGLLVGISSGAAAAAAIKLAQRPENAGKLFVAIFPSF<br>GERYLSTVLFDATRKEAEAMTFEA | MAATSSSAFLLNPLTSRHRPFKYSPELSSLSLSSRKAAAFDVSSAAFTLKRQSRSDVVCKAVSIK<br>PEAGVEGLNIADNAAQLIGKTPMVYLNNVVKGCVASVAAKLEIMEPCCSVKDRIGYSMITDA<br>EEKGLITPGKSVLVESTSGNTGIGLAFIAASKGYKLILTMPASMSLERRVLLRAFGAELVLTEPA<br>KGMTGAIQKAEEILKKTPNSYMLQQFDNPANPKIHYETTGPEIWEDTRGKIDILVAGIGTGGT<br>ITGVGRFIKERKPELKVIGVEPTESAILSGGKPGPHKIQGIGAGFVPKNLDLAIVDEYIAISSEEAI<br>ETSKQLALQEGLLVGISSGAAAAAAIQVAKRPENAGKLIAVVFPSFGERYLSTQLFQSIREECE<br>QMQPEL | | |

## D

| | | | | |
|---|---|---|---|---|
| Accession → | P47998 | P47999 | Q43725 | F4K5T2 |
| Gene name → | OAS-TL A | OAS-TL B | OAS-TL C | DES1 |

| | | | | |
|---|---|---|---|---|
| Protein area in upper activity band | 2.085E8 | 4.849E7 | 8.029E7 | 6.982E5 |
| Protein area in the middle activity band | 2.562E8 | 4.110E8 | 1.112E8 | 4.338E7 |
| Protein area in lowest activity band | 3.813E9 | 1.581E8 | 1.337E7 | 5.785E7 |

* Analysis of peptide sequences was performed by employing Proteome Discoverer™ Software ver. 1.4.1.14 (Thermo Fisher Scientific Inc. https://www.thermofisher.com/order/catalog/product/IQLAAEGABSFAKJMAUH

**Supplemental Table S2.** List of gene primers used for quantitative real-time PCR.

| Transcript | Accession number | Primer sequence (5'->3') | PCR product (bp) |
|---|---|---|---|
| *OASTL A1* | AT4G14880 | GCCTCGAGAATTGCTAAAGATGTGA<br>GGCTCAATCAGCACACTCTCTCC | 215 |
| *OASTL B* | AT2G43750 | CAGAGCCGGAGTGATGTTGTGT<br>CCTATCCTTGACACTGCAACATGG | 201 |
| *OASTL C* | At3G59760 | GGAGTTTTCCCAATGAGCTGAGAC<br>TCCTTCTCCTCCGCTTCTGATTT | 170 |
| *DES1*<br>Upstream T-DNA insertion | At5G28030 | TGGATGGTTGTGTGGCTCGT<br>ACCTCTTGAAGCCCCGATGC | 197 |
| *DES1*<br>Downstream T-DNA insertion | At5G28030 | AGTCGCCGGTGTTGGAACTG<br>CCTTGGATCAAATGTGGACCTGG | 147 |
| *ACTIN 2* | At3G18780 | TTGTGCTGGATTCTGGTGATGG<br>CCGCTCTGCTGTTGTGGTG | 167 |
| *EF 1-α* | AT5G60390 | CAGGACATCGTGATTTCATCAAGAAC<br>TCCATCTTGTTACAACAGCAAATCATCT | 190 |
